# Hog1/p38 and ZAKα drive Shwachman-Diamond syndrome and provide targets to improve cell growth

**DOI:** 10.64898/2026.02.05.703873

**Authors:** Nozomu Kawashima, Neha Prasad, Frank Tedeschi, Hrishikesh M. Mehta, Noah Saito, Cameron Jones, Xin Chen, Anca Manuela Hristodor, Gao Zhou, Joseph Luna, Marco Cipolli, Valentino Bezzerri, Seth J. Corey

## Abstract

Shwachman-Diamond syndrome (SDS) is a ribosomopathy characterized by neutropenia, pancreatic insufficiency, skeletal defects, and predisposition to leukemia. Most cases result from biallelic *SBDS* mutations that impairing 80S ribosome and polysome assembly. In yeast lacking *SDO1* (the *SBDS* ortholog), growth slows dramatically and the p38 ortholog Hog1 signaling is elevated by multiple types of stress. SBDS-deficient HeLa cells exhibited reduced proliferation and slowed cell cycling. The p38 kinase was constitutively activated in *SBDS* mutants and SDS patient-derived blood cells. Because ZAKα detects ribosome dysfunction, its activation links ribosomal defects to stress kinase pathways in SDS. Suppressing p38α or its upstream activator ZAKα restored cell growth and reduced stress signaling. These findings reveal an evolutionarily conserved-independent mechanism via p38 drives SDS pathophysiology and identifies stress kinases as potential therapeutic targets for ribosomal dysfunction.

## Introduction

Shwachman-Diamond syndrome (SDS) is an inherited bone marrow failure syndrome (IBMFS) characterized by neutropenia, exocrine pancreatic insufficiency, and skeletal abnormalities (*1*). In approximately 30%, transformation to the myeloid malignancies of myelodysplastic syndrome (MDS) or acute myeloid leukemia (AML) occurs (*2, 3*). AML is the leading cause of death in individuals with SDS due to therapy-resistance and high relapse rates. Stem cell transplantation is curative in a minority of those with myeloid malignancies but is associated with a heightened risk of toxicity (*4-6*). More than 90% of SDS patients carry biallelic mutations in the Shwachman-Bodian-Diamond syndrome (*SBDS*) gene, resulting in markedly decreased protein. In addition, mutations in *DNAJC21* (a member of the DnaJ Hsp40 family), *SRP54* (signal recognition particle 54), and *EFL1* (elongation factor-like 1 GTPase) have been linked to SDS or SDS-like syndromes (*7*). SBDS, EFL1, and DNAJC21 are evolutionally conserved and play critical roles in ribosome biogenesis.

SBDS plays a critical role in late ribosome maturation. It physically interacts with the GTPase EFL1 to displace eukaryotic translation initiation factor 6 (eIF6) from the 60S ribosomal large subunit and facilitates its assembly with the 40S ribosomal small subunit to form the mature 80S ribosome in the cytoplasm. Release of eIF6 is essential for the assembly of the 80S ribosome and polysomes (*8, 9*). Pathogenic genetic variants in *SBDS* and *EFL1* lead to decreased 80S formation and accumulation of 60S ribosomal large subunit and eIF6. SBDS deficiency and ribosomal dysfunction lead to activation of TP53 and TP53-dependent nucleolar surveillance pathway. Activation of the TP53 pathway is also observed in other IBMFS, such as Fanconi anemia and Diamond-Blackfan anemia (*1*). In SDS, selective pressure to bypass TP53-mediated cell cycle arrest and apoptosis drives the emergence of somatic mutations in *TP53* and *EIF6* (*10, 11*). While *EIF6* mutations or haploinsufficiency due to chromosomal deletion are not leukemogenic, *TP53* mutations follow a time-dependent progression from monoallelic to biallelic changes and are related to transformation to MDS/AML (*10-12*). These clinical findings suggest that acquisition of a loss-of-function *TP53* mutation may be maladaptive, whereas *EIF6* mutations may be adaptive and constitute somatic gene rescue. Using zebrafish lines with deletions of *sbds* and/or *eif6*, we confirmed somatic genetic rescue and found it correlated with modest reduction of Tp53 activation(*13*). Additionally, we reported that a loss-of-function mutation in *tp53* failed to rescue neutropenia or larval lethality in *sbds*^-/-^ zebrafish (*14*). RPL5 and RPL11 levels, which are increased in Diamond-Blackfan anemia and sequester MDM2 resulting in excessive TP53, are decreased in *sbds*^-/-^ zebrafish model (*14*). Thus, activation of the TP53 pathway appears to be necessary but not sufficient for the SDS phenotype downstream of the SBDS mutations.

Other signaling pathways have been suggested for the pathogenesis of SDS. Knockdown of *SBDS* mediated by short hairpin RNA induced greater release of reactive oxygen species (ROS), FAS-mediated apoptosis, and decreased cell proliferation in HeLa and the human myeloid leukemia TF-1 cell lines (*15, 16*). SBDS knockdown in HEK293 cells resulted in hypersensitivity to etoposide and ultraviolet-C radiation (*17*). Cre-mediated deletion of *Sbds* in a mouse mesenchymal stem cells shows mitochondrial dysfunction with increased ROS and DNA damage shown by phosphorylated H2AX histone (γH2AX) (*18*). Pancreas-specific Ptf1aCre–based *Sbds* conditional knockout in mouse pancreas upregulated *Tp53, Tgfb1*, and *Cdknb1* together with senescence-associated β-galactosidase in acinar cells. TP53 ablation partially rescued acinar cell function in this model (*19*). In SDS patient-derived samples, serum levels of TGFβ were elevated, and TGFβ signaling was upregulated in CD34^+^ bone marrow cells. TGF inhibitors (AVID200 and SD208) improved *in vitro* colony formation using patient-derived hematopoietic stem cells (*20*). Elevated levels of pro-inflammatory chemokines that are associated with the NF-κB pathway, such as interleukin-8, and the pathway, including CCL16 and CCL21 were reported in SDS patients (*21*). Signal transducer and activator of transcription 3 (STAT3) is hyperphosphorylated, and mechanistic target of rapamycin (mTOR) pathway was upregulated in SDS patient-derived cells (*22, 23*). All of those pathways do not directly emanate from a dysfunctional ribosome, prompting us to search for alternative mechanism(s) directly induced by impaired ribosomal maturation factors.

## Results

Because *S. cerevisiae* does not express TP53 or other cyclin-dependent kinase inhibitors, we studied *sdo1*Δ, the deletional mutant of *SDO1*, the yeast ortholog for *SBDS*, to identify TP53-independent stress pathway(s). In yeast lacking *SDO1*, demonstrated a slow growth phenotype (**Fig. 1**) with apparent doubling times for wild-type yeast 1.5 hr and *sdo1*Δ 9 hr (**fig. S1**). Mitogen-activated protein kinases, particularly Hog1, are activated by a variety of stress stimuli such as high osmotic load, oxidative or thermal stress, and nutritional depletion (*24*). Upon exposure to 0.8 M NaCl, Hog1 activation was both stronger and more prolonged in *sdo1*Δ cells compared to wild-type yeast (**Fig. 1B**). Upon activation, Hog1 translocates to the nucleus and increases the expression of genes, such as glycerol-3-phosphate dehydrogenase (GPD1). Consistent with Hog1 activation, there was marked increase in GPD1 transcript levels (**Fig. 1C**).

**Fig. 1.**
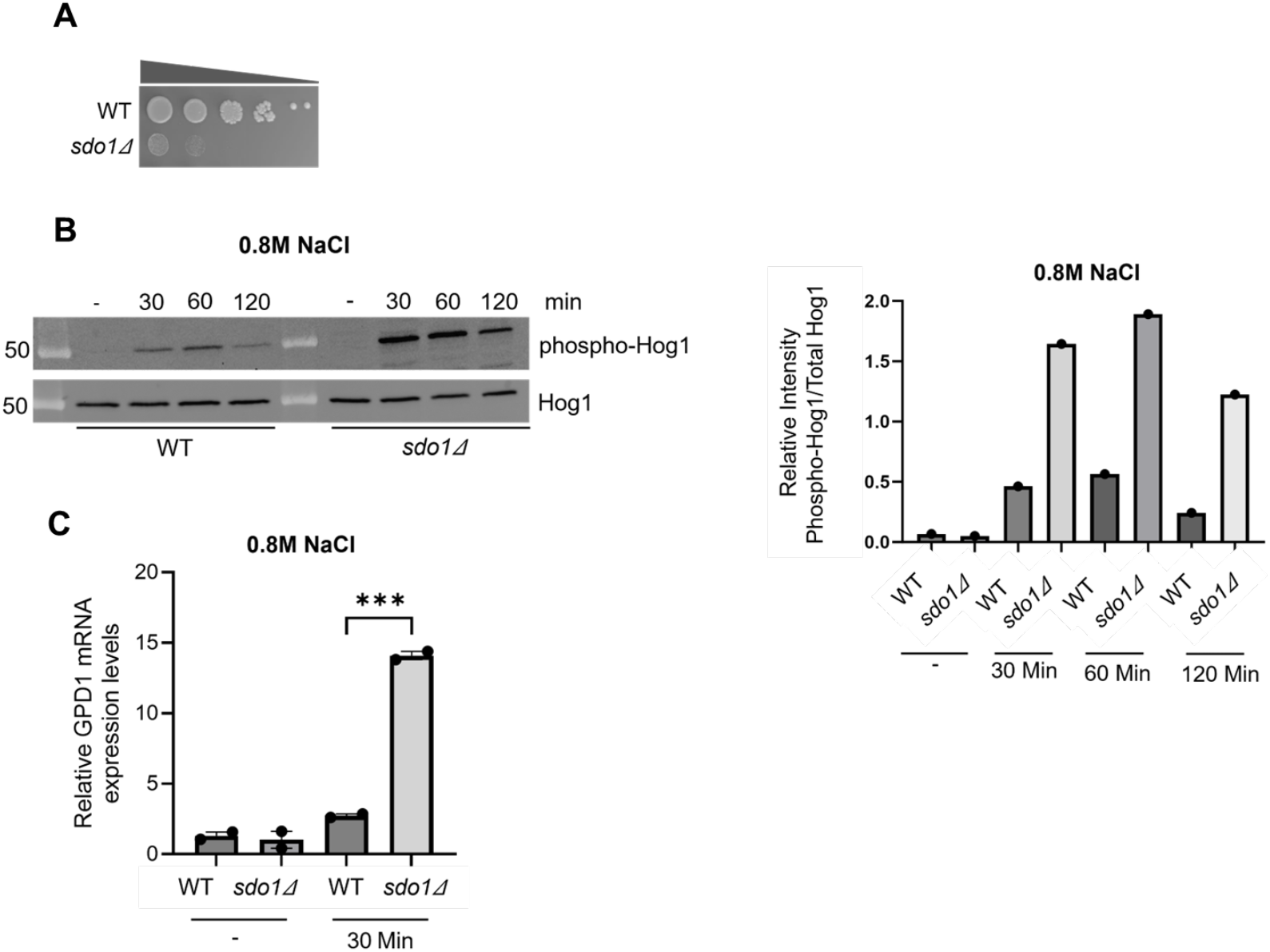
Reduced growth and activation of the stress-activated protein kinase Hog1. (**A**) Dilution drop assay of growth with ten-fold serial dilution of an equal number of WT and *sdo1*Δ cells were spotted on YPD media and incubated at 30°C for 5 days. Doubling times are 1.5 and 9 hr, respectively (**fig. S1**). (**B**) Lysates were prepared from WT and *sdo1*Δ cells, which were grown in YPD containing 0.8M NaCl and blotted for anti-phospho-Hog1. Blots were stripped and reprobed for total Hog1. Hog1 phosphorylation was greater and more sustained in *sdo1*Δ cells. The intensity of phosphorylated Hog1 was quantified using Image J software and normalized with respect to total Hog1p levels and shown in the form of bar diagram. (**C**) Quantitative PCR was performed for GPD1 on cDNA prepared from mRNA harvested from yeast grown in YPD medium containing 0.8 M NaCl. GPD1 upregulation was markedly elevated in the *sdo1*Δ cells. ***p<0.001

To study TP53-independent pathways in human cells, we used HeLa cells which are infected with human papillomavirus and carry its integrated genes, whose E6 protein interacts with the E3 ubiquitin ligase E6AP to degrade the TP53, rendering it functionally inactive despite the presence of a wild-type *TP53* (*25*). To study the role of SBDS deficiency in human cells, we mutagenized *SBDS* in HeLa cells by CRISPR/Cas9 editing (schema in **Fig. 2A**). Following single cell isolation, several clones were established. Clones were screened by immunoblotting for SBDS, and two clones were selected for decrease or loss of SBDS protein levels. Genotypes were confirmed by PCR amplicon deep sequencing (**fig. S2**). Clone S22 possessed a heterozygous mutation (NM_016038.4:c.188_198delGTCAGGTTGCC;p.G63Efs*13) and showed significantly decreased SBDS protein (thereafter, SBDS-KD). Clone S30 harbored homozygous mutations (NM_016038.4:c.201delA;p.K68Rfs*21) and showed only a trace of SBDS protein (thereafter, SBDS-KO; (**Fig. 2B**). Quantitative real-time PCR showed approximately 75% depletion of *SBDS* mRNA in SBDS-KD and 90% in SBDS-KO, possibly due to nonsense-mediated mRNA decay (**Fig 2C**). SBDS-KD and SBDS-KO were similar to parental cells (SBDS-WT) in morphology.

**Fig. 2.**
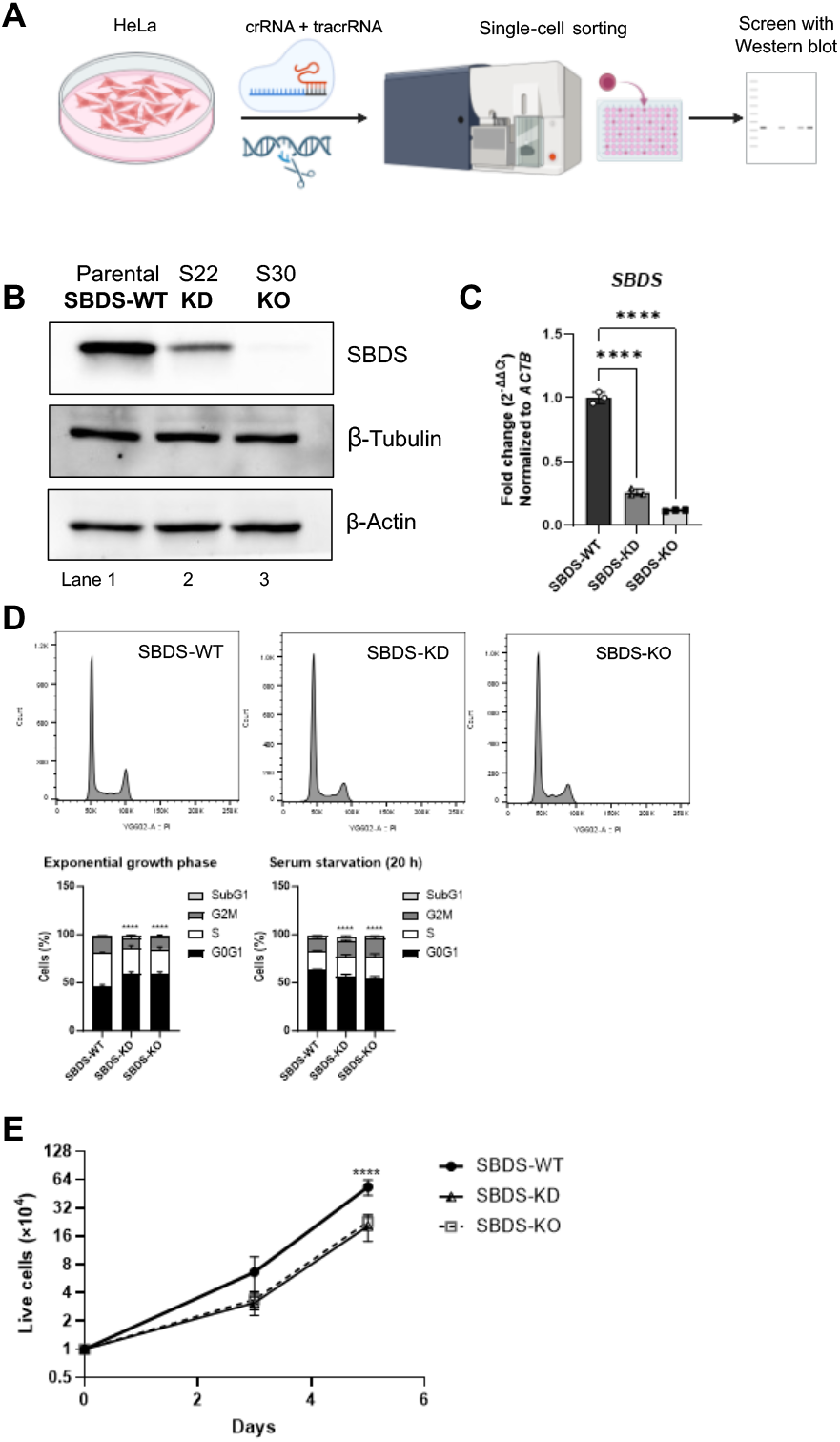
SBDS deficiency induced slow growth in yeast and HeLa cells. (**A**) Schema for single cell sorting of *SBDS*-edited HeLa cells. (**B**) Western blot for SBDS. Clone S22 showed a significantly decreased SBDS (thereafter, SBDS-KD) and Clone S30 showed no trace of SBDS (thereafter, SBDS-KO) compared to parental HeLa cells (SBDS-WT). (**C**) Quantitative real-time PCR showing nearly 75% depletion of *SBDS* mRNA in SBDS-KD and 90% in SBDS-KO compared to SBDS-WT, respectively. (**D**) Representative cell cycle analysis of unsynchronized cells in the exponential growth phase. The percentage of cells in G0-G1 was significantly increased in SBDS-KD and KO compared to WT, respectively. However, serum starvation for 20 h induced significantly higher percentages of G0-G1 in WT than in KD and KO, respectively. The chart shows mean of 3 biological replicates. (**E**) A decrease in the fold growth of SBDS-KO and KD cells compared to the WT cells 4 days after starting subculture. ****p<0.0001.

The percentages of cells in G0-G1 phase were significantly increased in SBDS-KD and KO compared to SBDS WT HeLa cells. However, when serum was depleted for 20 hours to induce G0-G1 arrest, those in G0-G1 phase were significantly decreased in SBDS-KD and KO than in SBDS-WT, respectively, which suggests a slowing in cell cycle progression of KD and KO (**Fig 2D**). Accordingly, a significant decrease in the growth of SBDS-KD and KO cells compared to the SBDS-WT was noted four days after starting subculture (**Fig 2E**).

To reveal the cellular vulnerability afforded by the loss of SBDS, we evaluated a variety of stressors, such as oxidative stress, nutritional deprivation, and genotoxic injury by gamma or ultraviolet radiation. Stress due to reactive oxygen species (ROS) demonstrated the most consistent stimulus. Organismal and cell line models indicate that eIF6 accumulation and activated TP53– CDKN1A cascade contribute considerably to SDS pathogenesis (*13, 26, 27*). While SBDS was depleted in SBDS-KO, the other ribosomal maturation factors, EFL1 and eIF6 were not significantly affected (**Fig. 3A**). Unexpectedly, CDKN1A (cyclin-dependent kinase inhibitor p21^Waf1^) was reduced 30 min after exposure to hydrogen peroxide in both SBDS-WT and SBDS-KO (**Fig. 3B**). These data suggest that eIF6 accumulation and activated TP53–CDKN1A pathway may not explain hypersensitivity of SBDS-KO to ROS. We evaluated key molecules for cell survival and apoptosis caused by exogenous stress. Total CDKN2A (p16), phospho-GSK3β, phospho-ERK1/2, phospho-AKT, phospho-CHK2, and phospho-γH2AX were not significantly different between SBDS-WT and SBDS-KO before and after exposure to ROS (**fig. S3**). Because ROS activates p38 to induce cell cycle arrest and apoptosis, we assessed its activation. When exposed to hydrogen peroxide, phosphorylation of p38 peaked 1 h after stimulation in SBDS-WT HeLa cells (**fig. S4**). Thus, we assessed phosphorylation of stress-activated protein kinases (SAPK) at this time point. Notably, basal levels of phospho-p38 and phospho-JNK were higher in SBDS-KO than in SBDS-WT (**Fig. 3C**,**D**). This observation was recapitulated in primary human peripheral blood mononuclear cells (PBMCs) from SDS patients carrying biallelic *SBDS* mutations (**table S1**). Compared with healthy controls, SDS patient-derived PBMCs exhibited a 10.7-fold increase in the phospho-p38/p38 ratio. (**Fig. 3E**). Since TGFβ signaling also activates p38 activation in a SMAD-independent mechanism,(*28*) our data aligns with reports of elevated TGFβ levels in patients (*20*).

**Fig. 3.**
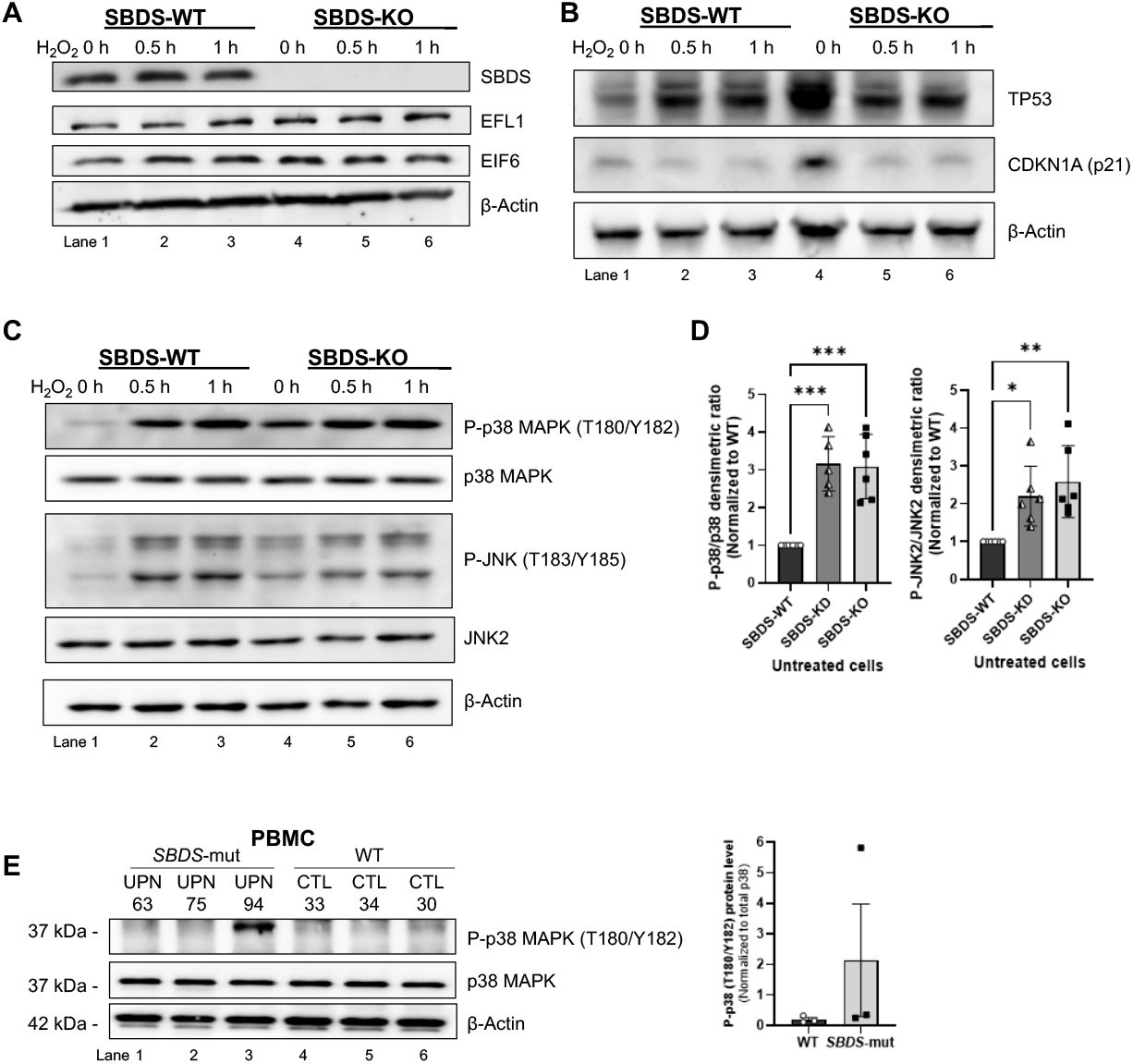
Protein and phosphorylation levels of SBDS-WT and KO HeLa cells post-exposure to hydrogen peroxide. (**A**) SBDS was depleted in SBDS-KO, whereas the other late ribosomal maturation factors, EFL1 and EIF6 were not affected. (**B**) CDKN1A (p21) was depleted 30 min after exposure to hydrogen peroxide (1 mM of H_2_O_2_) in both SBDS-WT and KO. (**C**) Basal levels of phospho-p38 MAPK and phospho-JNK were higher in SBDS-KO than in WT. Hydrogen peroxide induced upregulated phosphorylation in both WT and KO. (**D**) Densitometric analysis showing a significantly higher basal levels of phospho-p38 MAPK and phospho-JNK2 in SBDS-KD and KO than in WT. Values were expressed as mean of 5–6 biological replicates. (**E**) Phosphorylation levels in Shwachman-Diamond syndrome patient-derived samples. PBMCs from patients with Shwachman-Diamond syndrome showed a 10.7-fold increase in the phospho-p38/total p38 ratio compared to healthy controls. *p<0.05, ***p<0.001.

We tested whether Hog1/p38 inhibition could rescue their attenuated cell growth in SBDS/sdo1-deficient cells. To counter Hog1-mediated chronic stress and cell cycle arrest in yeast (*29*), the Ptc family of serine/threonine protein phosphatases inactivate Hog1 (*30*). In particular, Ptc1 catalyzes dephosphorylation of the threonine residue within Hog1’s activation loop (**fig. S5**). As shown in **Fig. 4A, right panel**, expression of Ptc1 rescues the growth of sdo1Δ yeast. As a control, we demonstrated that transduction of *sdo1Δ* yeast with a construct that expresses WT SDO1 corrected growth defect (**Fig. 4A, left panel**). Human p38 SAPK has four isoforms encoded by different genes. Of these, p38α is expressed at higher levels and mediate cell cycle arrest and/or apoptosis signaling, while p38β may have opposite effects (*31*). We focused on p38α and its specific inhibition. HeLa cells were treated with a potent p38α inhibitor, VX-745 (neflamapimod). VX-745 is 20 times more specific for p38α than p38β. Cell proliferation and live cells were significantly greater in VX-745-treated SBDS-deficient cells than nontreated cells (**Fig. 4B,C, fig. S6**). VX-745 did not affect SBDS-WT. Inhibition of p38α was confirmed by dephosphorylated MAP kinase-activated protein kinase 2 (MK2) in SBDS-KO which is regulated through direct phosphorylation by p38α (**Fig. 4D**). To confirm the findings of the p38α inhibitor, we transfected siRNA against *MAPK14* (encoding p38α isoform) in SBDS-WT, KD, and KO cells. Efficiency of gene silencing was assessed by quantitative real-time PCR (**fig. S7**). Western blot confirmed an siRNA-mediated decrease in the total p38 protein levels and particularly phospho-p38 in SBDS-KD and KO, respectively (**Fig. 4E**). We hypothesized that this suppression of phospho-p38 would rescue cell growth of the SBDS-deficient cells. The decreased cell proliferation in SBDS-KD and KO cells was partially corrected by siRNA against *MAPK14*. This siRNA had no significant effect on proliferation of SBDS-WT (**Fig. 4F**). Cell survival assessed by alamarBlue was also improved. Survival was significantly greater in SBDS-KD and KO cells at 3 and 5 days after transfection of siRNA against *MAPK14*, respectively, whereas WT was not significantly affected (**Fig. 4G**). Next, we asked whether p38 activation leading to slow cell growth could be rescued by radical scavengers. *N*-acetylcysteine or exogenous catalase treatment did not promote cell survival in SBDS-deficient cells (**fig. S8**). Accordingly, these scavengers did not significantly decrease spontaneous phosphorylation of p38 in SBDS-deficient cells **(fig. S9**). Altogether, our data suggest that Hog1/p38α activation reduces cell growth observed in *SBDS/sdo1-*null cells and that the genetic or chemical inhibition of these SAPK can rescue their growth arrest.

**Fig. 4.**
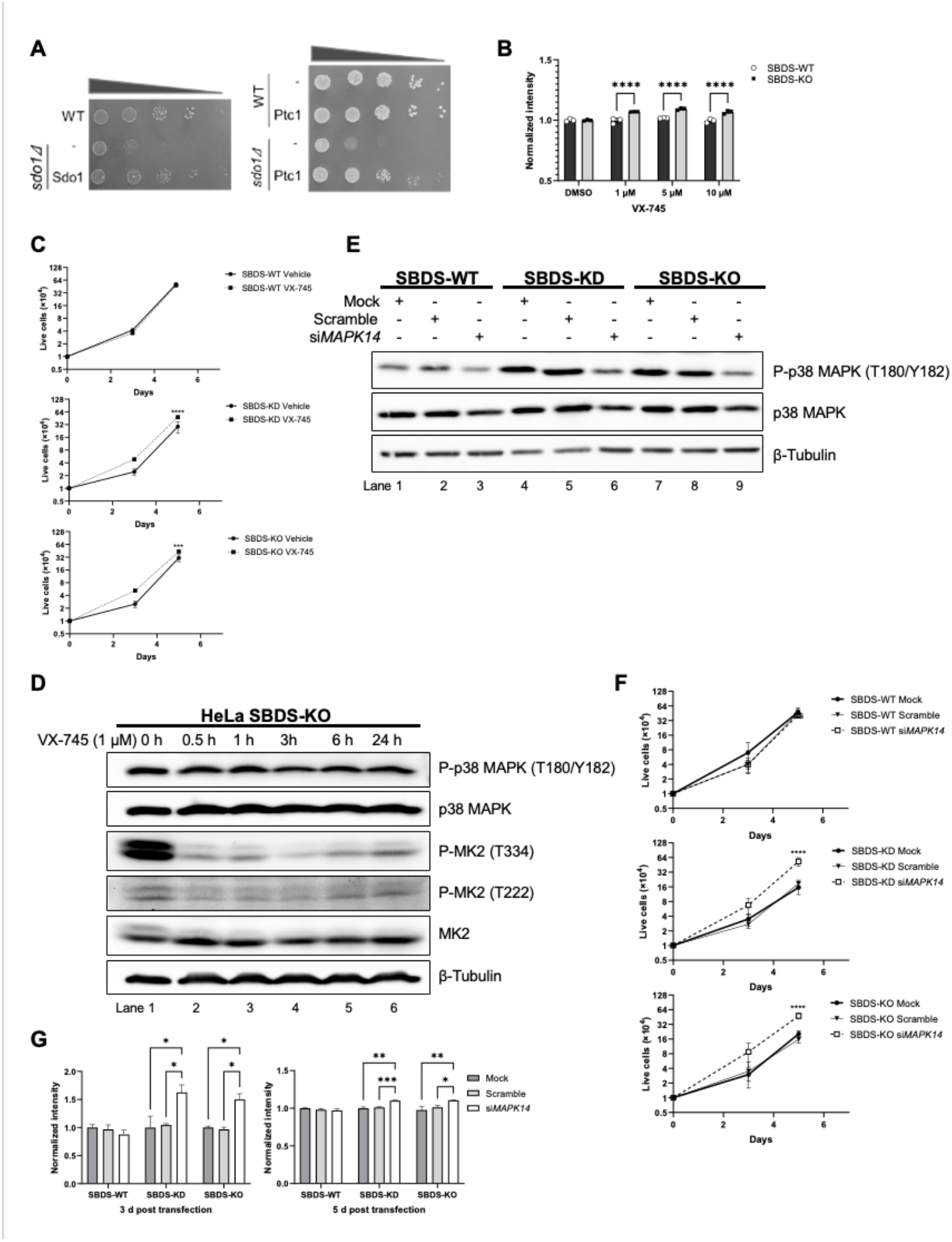
Genetic or chemical inhibition of Hog1/p38α activation rescued survival of SBDS depleted HeLa cells. **(A)** Dilution drop assay of wt, *sdo1*Δ, PTC1, and combinations thereof. (**B**) HeLa cells were treated with a potent p38α inhibitor, VX-745, for 5 days. Cell survival was assessed by alamarBlue assay. Live cells were significantly increased in SBDS-KO when exposed to ≥1 µM of VX-745. (**C**) Media containing either vehicle or VX-745 were replaced every 24 h for 5 days. A decrease in the fold growth of SBDS-KD and KO cells were rescued by VX-745. **(D)**MAP kinase-activated protein kinase 2 (MK2) is regulated through direct phosphorylation by p38α. VX-745 abrogated the spontaneous phosphorylation of MK2 (Thr334) in HeLa SBDS-KO. **(E)***MAPK14* (encoding p38α) siRNA were transfected into SBDS-WT, KD, and KO cells, respectively. Western blot showed decreased total p38 protein 5 days after siRNA transfection. Phospho-p38 were significantly decreased in SBDS-KD and KO. (**F**) A decrease in the fold growth of SBDS-KD and KO cells were rescued by siRNA against *MAPK14*, which had no significant effect on proliferation of SBDS-WT cells. (F) Cell survival assessed by alamarBlue assay was significantly more in SBDS-KD and KO 3 and 5 days after transfection of siRNA against *MAPK14*. *p<0.05, **p<0.01, ***p<0.001, ****p<0.0001.

The SAPK are activated in response to various stress stimuli in a cell-dependent manner, either directly or indirectly targeting downstream proteins to control cell cycle checkpoints. Recently, a member of the MAPK3 family, the long-splice isoform ZAKα encoded by *MAP3K20* binds to mature ribosomes and sense their stalling and/or collision caused by exogenous and endogenous damage in metazoans (*32-35*). Upon activation, ZAKα auto-phosphorylates and phosphorylates downstream substrates including p38 and JNK to exert cell cycle arrest. Because SBDS is associated with late ribosome maturation and its deficiency contributes to vulnerability to ROS as shown above, we hypothesized that spontaneously activated SAPK might be induced by activated ZAKα. ZAKα was immunoprecipitated from cell lysates and probed for pan-threonine phosphorylation. ZAKα was constitutively more phosphorylated in SBDS-KD and KO than in WT (**Fig. 5A**). Conversely, immunoprecipitation with anti-phospho-threonine antibodies showed co-precipitation of ZAKα in SBDS-KD and KO lysates, whereas ZAKα was nearly undetectable in SBDS-WT (**Fig. 5B**). We transfected siRNA against *MAP3K20* to successfully deplete ZAKα. Efficiency of *MAP3K20* silencing was assessed by quantitative real-time PCR (**fig. S7**). ZAKα depletion significantly decreased basal phosphorylation of p38 in SBDS-KD and KO to the level of WT, suggesting ZAKα as a major activator of p38 in SBDS insufficiency (**Fig. 5C**). Depletion of ZAKα due to *MAPK3K20* siRNA rescued reduced growth of SBDS-KO cells by 5 days post-transfection (**Fig. 5D**). Cell survival was also significantly improved in SBDS-KO cells transfected with *MAPK3K20* siRNA (**Fig. 5E**). Finally, we asked whether this role of ZAKα in p38 phosphorylation in SBDS deficiency be due to a direct result from attenuated ribosome maturation or due to high ROS burden induced by this ribosomal stress. As shown above, exogenous hydrogen peroxide activated p38 within 0.5 h (**Fig. 3C**). Accordingly, hydrogen peroxide activated ZAKα in SBDS-WT cells within 0.5 h, although a potent ROS scavenger, *N*-acetylcysteine, did not mitigate this phosphorylation (**fig. S10**). Neither did *N*-acetylcysteine nor exogenous catalase ameliorate auto-phosphorylation of ZAKα in SBDS-deficient cells (**fig. S11**), which is consistent with their failure to suppress activated p38 in SBDS-KD and KO (**fig. S9**). These data suggest that ROS reduction by antioxidants does not contribute to suppress the activated ZAKα/p38 pathway.

**Fig. 5.**
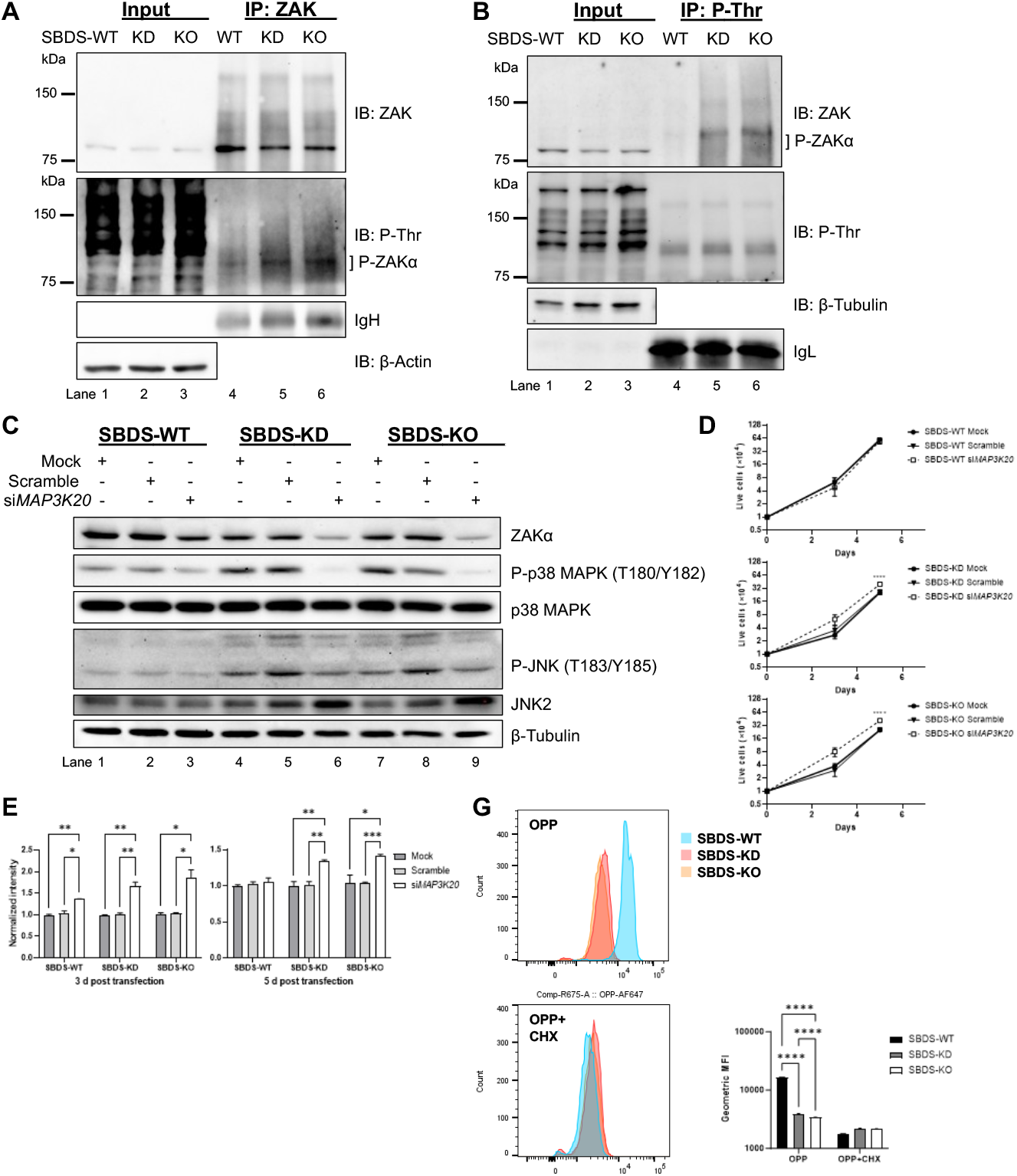
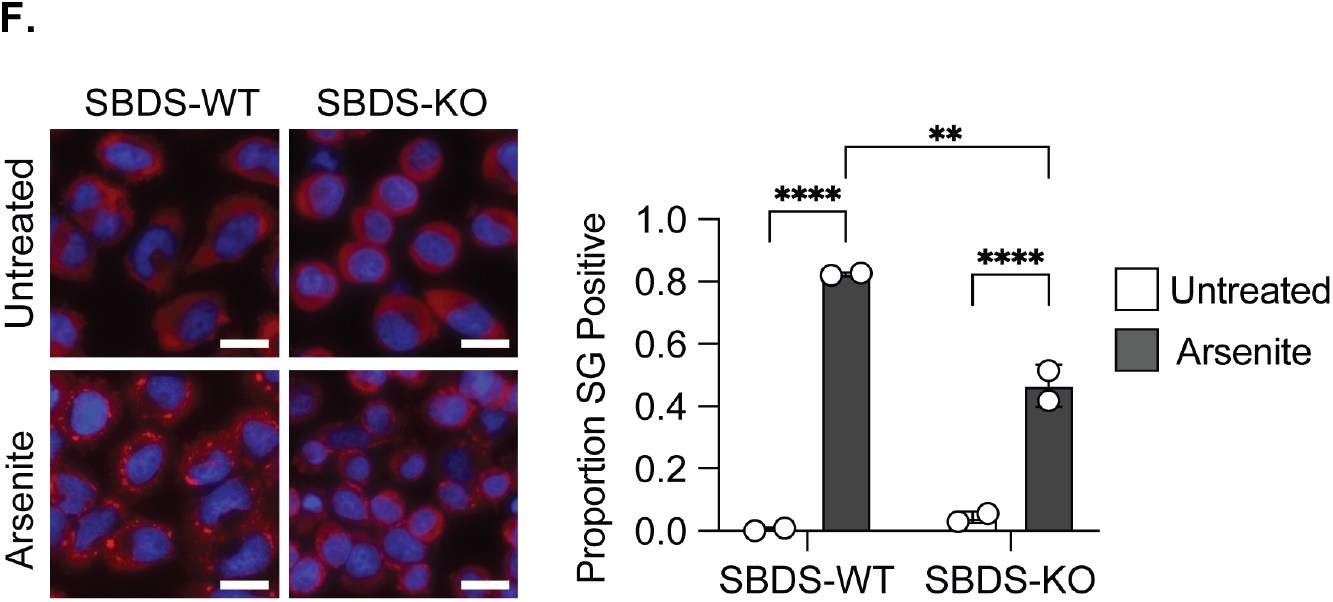
Ribosome stalling triggers spontaneous ZAKα activation in SBDS-deficient cells, and ZAKα depletion restores cell survival. (**A**) Immunoprecipitation (IP) with anti-ZAK antibodies followed by immunoblot (IB) showing hyperphosphorylated ZAKα in SBDS knockdown (KD) and knockout (KO) cells. (**B**) IP with anti-phospho-threonine antibodies and IB with anti-ZAK antibodies showing ZAKα co-precipitation in SBDS-KD and KO, but not SBDS-WT cells. (**C**) Western blot confirming ZAKα depletion 5 days after *MAP3K20* siRNA transfection; spontaneous phospho-p38 levels were significantly reduced in SBDS-KD and KO cells. (**D**) Cell culture growth curves showing reduced proliferation of SBDS-KO cells rescued by *MAP3K20* siRNA at day 5. **(E)**Cell survival significantly improved in SBDS-KO cells after *MAP3K20* siRNA transfection. **(F)**Basal and induced stress granule formation occurred in SBDS-KO cells and increased with sodium arsenite; cells were stained with a G3BP1 antibody with and without treatment of 500 uM sodium arensite for 45min. White scale bar 25um.; SBDS-WT cells formed more granules under oxidative stress, suggesting chronic stress adaptation in SBDS-KO. (**G**) O-propargyl-puromycin (OPP) assay showing decreased nascent protein synthesis in SBDS-KD and KO cells compared to WT; cycloheximide (CHX, 50 µg/mL) pretreatment inhibited OPP incorporation across all genotypes. MFI, mean fluorescence intensity. *p < 0.05, **p < 0.01, ****p < 0.0001.

Because deficiency of SBDS causes stoichiometric imbalance between the 40S ribosomal small subunit, 60S ribosomal large subunit that decreases 80S monosome formation, we hypothesize that a chronic ribotoxic stress state results. In addition to activating ZAKα/p38 pathway, stress granules are formed as a downstream adaptive translational checkpoint. We observed low level basal stress granule formation in SBDS-KO HeLa cells that increased with sodium arsenite treatment but was impaired relative to WT controls (**Fig. 5F**). These results are compatible with a chronic stress state and attempts to mitigate stress through adaptive mechanisms to be defined.(*36*) One result of chronic stress granule formation is diminished global protein translation. Not unexpectedly, naïve SBDS-WT HeLa cells displayed a greater formation in stress granules in response to oxidative stress. These results are compatible with a chronic stress state and attempts to mitigate stress through adaptive mechanisms to be defined. One result of chronic stress granule formation is diminished global protein translation. O-propargyl puromycin (OPP), a puromycin analog, is efficiently incorporated into newly translated proteins which is fluorescently labeled and detected by flow cytometry in a highly-specific and sensitive manner (*37, 38*). Strikingly, incubation with OPP for 30 min showed significantly decreased incorporation of OPP in SBDS-KD and KO compared to SBDS-WT, suggesting nascent protein synthesis is significantly mitigated in SBDS-deficient cells (**Fig. 5G**). Cycloheximide (CHX), a tRNA inhibitor, successfully depleted OPP incorporation in all cells. OPP or CHX treatment up to 60 min and 75 min did not cause cell death assessed by trypan blue stain in these cells, respectively.

Altogether, those data suggest that a role of ZAKα in stimulating the p38 phosphorylation in cells deficient in SBDS. ZAKα itself is activated as a direct result from ribosome dysfunction, causing ribosome stalling. While there is no yeast ortholog for ZAKα, Hog1 may serve as an effector for ribotoxic stress. To demonstrate ribosome stalling in *sdo1*Δ yeast, we analyzed polysome profiles, comparing WT with mutant strains. Polysome profiling demonstrated an increase in disome and trisome peaks in the *sdo1*Δ yeast (**Fig. 6A**). Using ORFik, a ribosome stalling analysis software (*39*), there was increased stalling in the *sdo1*Δ yeast compared to the WT strain (**Fig. 6B**). Ribosome profiling revealed a global increase in stalling scores and coverage variability in *sdo1*Δ cells, indicating widespread defects in elongation. Most expressed genes showed elevated ribosome stalling compared with WT, consistent with impaired ribosome dynamics as a primary driver of stress pathway activation (**lower right panel in Fig. 6B**).

**Fig. 6.**
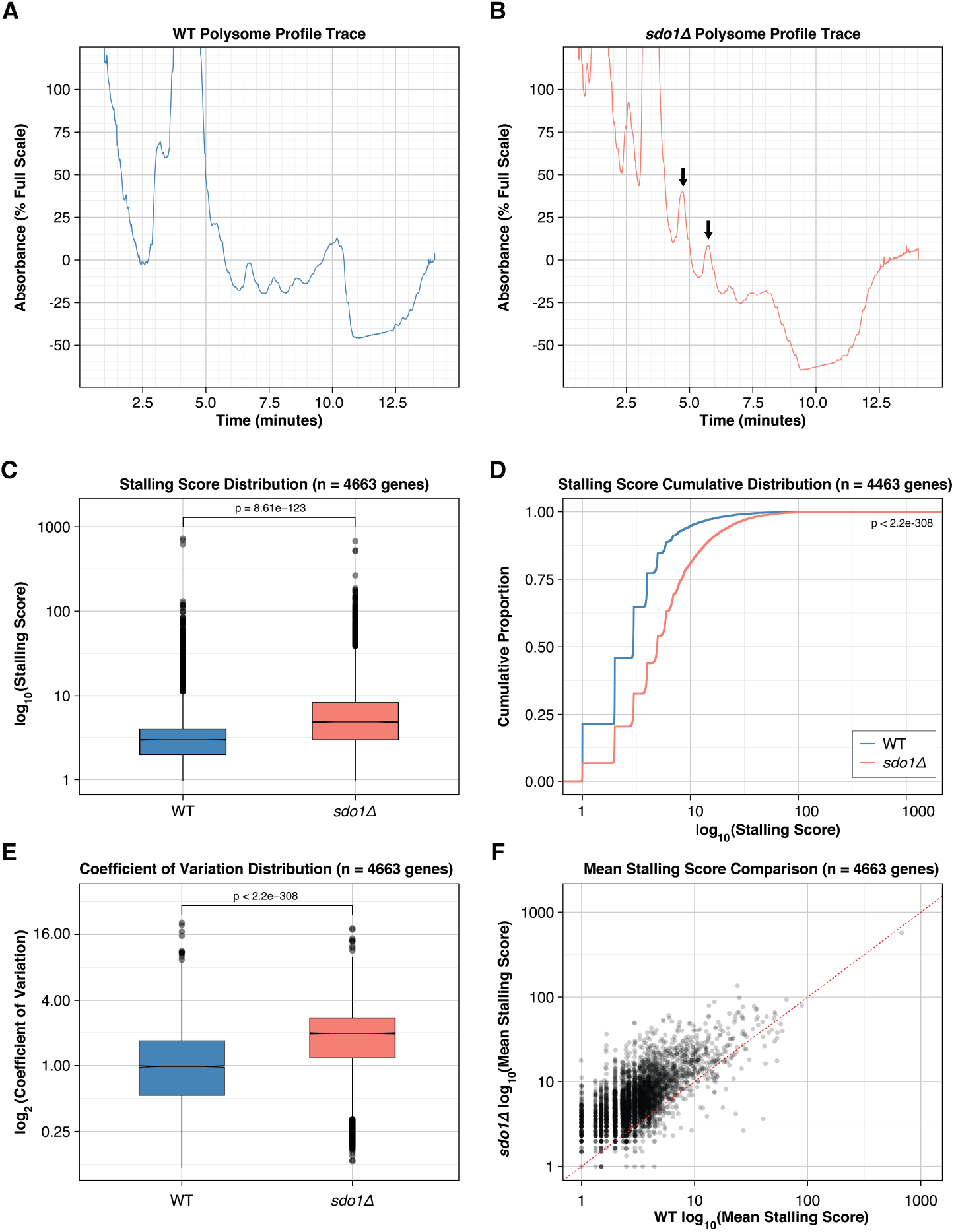
Polysome and Ribosome Analysis in SBDS mutants. **(A–B)** Representative polysome profile traces from yeast **WT** (A) and **sdo1Δ** (B) cells following sucrose gradient fractionation. UV absorbance at 254 nm is plotted as a function of time. Arrows in (B) mark ribosome disome and trisome peaks. **(C)** Boxplots showing the distribution of gene-level stalling scores for WT and *sdo1Δ* across 4,663 mRNA-filtered genes. Stalling score is calculated as log_10_(Max ribosome footprint coverage / Mean coverage) per transcript. Two-sided t-test p = 8.61 × 10^−123^. **(D)** Cumulative distribution of stalling scores for WT and *sdo1Δ* genes. The plot shows the cumulative proportion of genes at or below each stalling score value (log_10_ scale). Kolmogorov– Smirnov test p < 2.2 × 10^−308^. **(E)** Boxplots of the coefficient of variation (CV) of ribosome footprint coverage per gene, calculated as log_2_(SD/Mean), for WT and *sdo1Δ* (n = 4,663 genes). Two-sided t-test p < 2.2 × 10^−308^. **(F)** Scatter plot comparing mean stalling scores for individual genes in WT (x-axis) and *sdo1Δ* (y-axis), both shown on a log_10_ scale. Each point represents one gene.

## Discussion

We have demonstrated activation of the Hog1/p38 SAPK in yeast and human cells lacking SBDS/SDO1 and that genetic or chemical inhibition of Hog1/p38 can rescue their reduced cell growth. Thus, there are evolutionarily conserved SBDS-EFL1-eIF6 and Hog1/p38 pathways in normal and dysfunctional eukaryotic ribosome assembly. In higher eukaryotic cells, there are additional mediators, e.g., ZAKα and TP53, to manage ribosomal stress.

When ribosomal stress is chronic (as in germline *SBDS* mutations), loss-of-function mutations in TP53 occurs and myeloid malignancy arises. Sequencing of SDS patients across the lifespan reveals time-dependent acquisition of *TP53* mutations (*10, 11*). The lifetime risk of developing myeloid malignancy (MDS/AML) may be as high as 30-40% (*40*). In SDS patients undergoing anti-leukemic therapy, there is increased sensitivity to chemotherapy side effects concomitant with chemoresistance of myeloid cells in patients with SDS and AML (*2, 4, 41*). We found that SBDS-deficient cells exhibited selective sensitivity to cell stressors, with inducers of oxidative stress being the most consequential. However, exogenous ROS promptly induced depletion of CDKN1A, which also challenges a necessary and sufficient role for TP53 and its mediators in SBDS deficiency. Instead, cell stressors with selective sensitivity to SBDS deficiency recruited SAPK activation, emphasizing its role for SDS.

Transient senescence can restrain tumor growth and support tissue regeneration, thereby exerting beneficial effects. Myelosuppressive interferons and other cytokines can activate the p38 MAPK pathway, which may in turn amplify NF-κB- or C/EBPβ-dependent inflammation, fostering a self-reinforcing feedback loop leading to TP53-mediated cell cycle arrest (*42*). Bone marrow cells from patients with acquired aplastic anemia and MDS display activated p38 and that p38α inhibitor rescued hematopoietic cells for self-renewal and proliferation (*43*). Most IBMFS, including Fanconi anemia and Diamond-Blackfan anemia, have already been linked to accelerated senescence and chronic systemic inflammation. We have recently demonstrated that SDS patients display constitutive systemic inflammation dominated by senescence-associated secretory phenotype mediators such as IL-6 and IL-8 (*44*). Persistent senescence, such as that triggered by sublethal genetic damages, including deficiency of SBDS, is detrimental and promotes chronic inflammation and tumor initiation, especially if released from the control mechanism(*45*). What was an adaptive response becomes a maladaptive, leukemogenic one with biallelic loss of wild-type *TP53*(*12*).

Upstream of p38α lies ZAKα, which mediates ribotoxic stress response that occurs upon ribosome stalling and collisions. Depleting ZAKα induced less phosphorylation of p38 in SBDS-deficient cells and cell proliferation improved which was consistent with results with p38α inhibition. Protein synthesis is tightly regulated, and its abrupt stop due to ribosome stalling and/or collisions is promptly sensitized and rescued (*46*). Ribosome stalling and collisions can be induced by local translocation-blocking mRNA lesion induced by e.g., UV, elongation distress (anisomycin, emetine, and sarcin/ricin). Ribosome stalling can be induced by general cell stress such as the amino acid depletion/starvation and the ROS which was recently identified to cause directly ribosome stalling and activate ZAKα (*47-49*). Additionally, RNA viral infection can induce ZAKα activation (*34*) and this may be mediated by cellular antiviral mechanism of RNase L to digest viral or cellular RNA (*50*). ZAKα binds directly to the translationally active ribosomes and senses ribosome stalling and/or collisions via partially redundant domains.(*33*) It auto-phosphorylates and this is associated with ribosome scanning, relieve the SAM-domain-mediated auto-inhibition, and phosphorylation of downstream MAP2Ks 3/6, 4/7, and subsequently, SAPK (*33, 50*). This series of activating SAPK constitutes ribotoxic stress reaction. Interestingly, ZAKα is activated during erythroid differentiation of human leukemia cell line K562, inducing the NLRP1 inflammasome-mediated GATA1 degradation. Multikinase inhibitor nilotinib reversed this and alleviated neutrophilia in the *spint1a* mutated zebrafish model of neutrophilic inflammation (*51*). We showed that reduced protein synthesis in SBDS-deficient cells, which will directly activate ZAKα. Although it still requires additional evidence to determine whether steady activation of ZAKα is due to direct stress by “slow” translation caused by defects in ribosomal maturating factors or an indirect result from endogenous ROS or other ribotoxic stressors, our data implicated a new mechanism for activation of ZAKα/SAPK in SDS pathogenesis.

Our studies support the existence of a Hog1/p38 and ZAKα activation/growth arrest pathway, independent of TP53, in SDS. Chronic stimulation of p38 might then lead to neoplastic transformation via additional pressures on TP53 pathways. In this scenario, targeting stress-activated kinases with ZAKα or p38α inhibitors, or employing senomorphics such as natural antioxidants and anti-inflammatory agents, emerges as a promising strategy to halt progression toward MDS/AML, reduce the toxicity associated with chemotherapy, and alleviate additional pathological manifestations of SDS.

## Acknowledgments

The authors acknowledge the assistance of the Case Western Reserve University School of Medicine Light Microscopy Imaging Facility and support by NIH grant #S10OD02499601.

## Funding

SENSHIN Medical Research Foundation (NK) Mochida Memorial Foundation (NK)

Japan Society for the Promotion of Science (NK) NIH R01DK128173 (SJC)

NIH R21CA159203 (SJC)

DOD Idea Award (SJC) Hyundai Hope on Wheels (SJC) VeloSano (NK, SJC)

Lisa Dean Moseley Foundation Award (SJC)

## Author contributions

Funding acquisition: SJC, NK, VB, MC

Designed experiments: NK, NP, FT, RMD, FT, VB, MC, SJC

Performed experiments: NK, NP, FT, HMM, NS, CJ, XC, AMH, GJ, VB Analyzed the data: NK, NP, HM, FT, JL, VB, SJC

Wrote the manuscript: NK, NP, FT, HMM, JL, VB, SJC

## Data, code, and materials availability

Available from the corresponding author

## References and Notes

1. N. Kawashima, V. Bezzerri, S. J. Corey, The Molecular and Genetic Mechanisms of Inherited Bone Marrow Failure Syndromes: The Role of Inflammatory Cytokines in Their Pathogenesis. Biomolecules 13, 1249–1268 (2023).

2. S. Cesaro et al., A Prospective Study of Hematologic Complications and Long-Term Survival of Italian Patients Affected by Shwachman-Diamond Syndrome. J Pediatr 219, 196–201 e191 (2020).

3. E. Furutani et al., Hematologic complications with age in Shwachman-Diamond syndrome. Blood Adv 6, 297–306 (2022).

4. S. Cesaro et al., Stem Cell Transplantation in Patients Affected by Shwachman-Diamond Syndrome: Expert Consensus and Recommendations From the EBMT Severe Aplastic Anaemia Working Party. Transplant Cell Ther 28, 637–649 (2022).

5. J. Donadieu et al., Hematopoietic stem cell transplantation for Shwachman-Diamond syndrome: experience of the French neutropenia registry. Bone Marrow Transplant 36, 787–792 (2005).

6. K. Myers et al., Hematopoietic Stem Cell Transplantation for Shwachman-Diamond Syndrome. Biol Blood Marrow Transplant 26, 1446–1451 (2020).

7. N. Kawashima, U. Oyarbide, M. Cipolli, V. Bezzerri, S. J. Corey, Shwachman-Diamond syndromes: clinical, genetic, and biochemical insights from the rare variants. Haematologica 108, 2594–2605 (2023).

8. T. F. Menne et al., The Shwachman-Bodian-Diamond syndrome protein mediates translational activation of ribosomes in yeast. Nature Genetics 39, 486–495 (2007).

9. F. Weis et al., Mechanism of eIF6 release from the nascent 60S ribosomal subunit. Nat Struct Mol Biol 22, 914–919 (2015).

10. S. Tan et al., Somatic genetic rescue of a germline ribosome assembly defect. Nat Commun 12, 5044 (2021).

11. A. L. Kennedy et al., Distinct genetic pathways define pre-malignant versus compensatory clonal hematopoiesis in Shwachman-Diamond syndrome. Nat Commun 12, 1334 (2021).

12. H. E. Machado et al., Convergent somatic evolution commences in utero in a germline ribosomopathy. Nat Commun 14, 5092 (2023).

13. U. Oyarbide et al., Reduced EIF6 dosage attenuates TP53 activation in models of Shwachman-Diamond syndrome. J Clin Invest 135, e187778 (2025).

14. U. Oyarbide et al., SBDS(R126T) rescues survival of sbds (-/-) zebrafish in a dose-dependent manner independently of Tp53. Life Sci Alliance 6, e202201856 (2023).

15. C. Ambekar, B. Das, H. Yeger, Y. Dror, SBDS-deficiency results in deregulation of reactive oxygen species leading to increased cell death and decreased cell growth. Pediatr Blood Cancer 55, 1138–1144 (2010).

16. K. Watanabe et al., SBDS-deficiency results in specific hypersensitivity to Fas stimulation and accumulation of Fas at the plasma membrane. Apoptosis 14, 77–89 (2009).

17. H. L. Ball et al., Shwachman-Bodian Diamond syndrome is a multi-functional protein implicated in cellular stress responses. Hum Mol Genet 18, 3684–3695 (2009).

18. M. H. G. P. Raaijmakers et al., Bone progenitor dysfunction induces myelodysplasia and secondary leukaemia. Nature 464, 852–857 (2010).

19. M. E. Tourlakis et al., In Vivo Senescence in the Sbds-Deficient Murine Pancreas: Cell-Type Specific Consequences of Translation Insufficiency. PLOS Genetics 11, e1005288 (2015).

20. C. E. Joyce et al., TGFbeta signaling underlies hematopoietic dysfunction and bone marrow failure in Shwachman-Diamond Syndrome. J Clin Invest 129, 3821–3826 (2019).

21. E. Furutani et al., Inflammatory manifestations in patients with Shwachman–Diamond syndrome: A novel phenotype. American Journal of Medical Genetics. Part A 182, 1754–1760 (2020).

22. V. Bezzerri et al., New insights into the Shwachman-Diamond Syndrome-related haematological disorder: hyper-activation of mTOR and STAT3 in leukocytes. Sci Rep 6, 33165 (2016).

23. A. Vella et al., mTOR and STAT3 Pathway Hyper-Activation is Associated with Elevated Interleukin-6 Levels in Patients with Shwachman-Diamond Syndrome: Further Evidence of Lymphoid Lineage Impairment. Cancers (Basel) 12, 597–616 (2020).

24. G. Du et al., The relationship mammalian p38 with human health and its homolog Hog1 in response to environmental stresses in Saccharomyces cerevisiae. Front Cell Dev Biol 13, 1522294 (2025).

25. J. C. K. Wang et al., Structure of the p53 degradation complex from HPV16. Nat Commun 15, 1842 (2024).

26. P. Rujkijyanont, S. L. Adams, J. Beyene, Y. Dror, Bone marrow cells from patients with Shwachman-Diamond syndrome abnormally express genes involved in ribosome biogenesis and RNA processing. Br J Haematol 145, 806–815 (2009).

27. U. Oyarbide et al., Loss of Sbds in zebrafish leads to neutropenia and pancreas and liver atrophy. JCI Insight 5, e134309 (2020).

28. C. H. Heldin, A. Moustakas, Signaling Receptors for TGF-beta Family Members. Cold Spring Harb Perspect Biol 8, a022053 (2016).

29. X. Escote, M. Zapater, J. Clotet, F. Posas, Hog1 mediates cell-cycle arrest in G1 phase by the dual targeting of Sic1. Nat Cell Biol 6, 997–1002 (2004).

30. J. Warmka, J. Hanneman, J. Lee, D. Amin, I. Ota, Ptc1, a type 2C Ser/Thr phosphatase, inactivates the HOG pathway by dephosphorylating the mitogen-activated protein kinase Hog1. Mol Cell Biol 21, 51–60 (2001).

31. B. Canovas, A. R. Nebreda, Diversity and versatility of p38 kinase signalling in health and disease. Nat Rev Mol Cell Biol 22, 346–366 (2021).

32. C. C. Wu, A. Peterson, B. Zinshteyn, S. Regot, R. Green, Ribosome Collisions Trigger General Stress Responses to Regulate Cell Fate. Cell 182, 404–416.e414 (2020).

33. A. C. Vind et al., ZAKα Recognizes Stalled Ribosomes through Partially Redundant Sensor Domains. Mol Cell 78, 700–713.e707 (2020).

34. K. S. Robinson et al., ZAKα-driven ribotoxic stress response activates the human NLRP1 inflammasome. Science 377, 328–335 (2022).

35. G. Snieckute et al., Ribosome stalling is a signal for metabolic regulation by the ribotoxic stress response. Cell Metab 34, 2036–2046.e2038 (2022).

36. Y. Adachi et al., Chronic stress antagonizes formation of stress granules. iScience 29, 114556 (2026).

37. L. Hidalgo San Jose, R. A. J. Signer, Cell-type-specific quantification of protein synthesis in vivo. Nat Protoc 14, 441–460 (2019).

38. J. C. Hsu, J. B. Pawlak, M. Laurent-Rolle, P. Cresswell, Protocol for assessing translational regulation in mammalian cell lines by OP-Puro labeling. STAR Protoc 3, 101654 (2022).

39. H. Tjeldnes et al., ORFik: a comprehensive R toolkit for the analysis of translation. BMC Bioinformatics 22, 336 (2021).

40. J. Donadieu et al., Classification of and risk factors for hematologic complications in a French national cohort of 102 patients with Shwachman-Diamond syndrome. Haematologica 97, 1312–1319 (2012).

41. Z. Hudda, K. C. Myers, Posttransplant complications in patients with marrow failure syndromes: are we improving long-term outcomes? Hematology Am Soc Hematol Educ Program 2023, 141–148 (2023).

42. A. Hernandez-Segura, J. Nehme, M. Demaria, Hallmarks of Cellular Senescence. Trends Cell Biol 28, 436–453 (2018).

43. T. A. Navas et al., Inhibition of overactivated p38 MAPK can restore hematopoiesis in myelodysplastic syndrome progenitors. Blood 108, 4170–4177 (2006).

44. G. Sabbioni et al., Constitutive systemic inflammation in Shwachman-Diamond Syndrome. Mol Med 31, 81 (2025).

45. M. Milanovic et al., Senescence-associated reprogramming promotes cancer stemness. Nature 553, 96–100 (2018).

46. M. C. J. Yip, S. Shao, Detecting and Rescuing Stalled Ribosomes. Trends Biochem Sci 46, 731–743 (2021).

47. C. C. Wu, A. Peterson, B. Zinshteyn, S. Regot, R. Green, Ribosome Collisions Trigger General Stress Responses to Regulate Cell Fate. Cell 182, 404–416 e414 (2020).

48. G. Snieckute et al., ROS-induced ribosome impairment underlies ZAKalpha-mediated metabolic decline in obesity and aging. Science 382, eadf3208 (2023).

49. K. S. Robinson et al., ZAKalpha-driven ribotoxic stress response activates the human NLRP1 inflammasome. Science 377, 328–335 (2022).

50. J. Xi et al., Initiation of a ZAKalpha-dependent ribotoxic stress response by the innate immunity endoribonuclease RNase L. Cell Rep 43, 113998 (2024).

51. L. Rodríguez-Ruiz et al., ZAKα/P38 kinase signaling pathway regulates hematopoiesis by activating the NLRP1 inflammasome. EMBO Mol Med 15, e18142 (2023).

