## Supplemental information for "Hog1/p38 and ZAKα drive Shwachman-Diamond syndrome and provide targets to improve cell growth"

[1]Departments of Pediatrics and Heart, Lung, and Blood, Cleveland Clinic, Cleveland, OH; [2]Center for RNA Science and Therapeutics; Case Western Reserve University, Cleveland, OH; [3]Deprtment of Biochemistry, Case Western Reserve University, Cleveland, OH; [4]Cystic Fibrosis Center, Azienda Ospedaliera Universitaria Integrata, Verona, Italy; [5]Department of Health and Health Professions, Link Campus University, Rome, Italy

†These authors contributed equally to this work.

†These authors contributed equally to this work.

**The PDF file includes:**

1. Materials and Methods
2. Figs. S1 to S11
3. Tables S1 to S4

Supplementary Text: Materials and Methods

Figs. S1 to S11

Tables S1 to S4

References 3

Movies: N/A

Audio: N/A

Data: N/A

**Materials and Methods**

*Cells and reagents*. The human cervical cancer cell line HeLa cells were purchased from ATCC and maintained in DMEM supplemented with 10% fetal bovine serum and 1% penicillin/streptomycin. For serum starvation, fetal bovine serum was substituted with 0.1% bovine serum albumin. Sources for reagents to stress cells are listed in the **Supplemental Table 2**. Yeast strains (*SDO1* and *sdo1*Δ) used in this study were obtained from EUROSCARF and belong to the W303 strain. *sdo1Δ*::KanMX4 haploid strain was constructed by sporulating heterozygous W303 (W303, SDO1/ *sdo1Δ*::KanMX4 strain). A haploid strain harboring the kanMX4 cassette was selected using G418 (200 µg/ml) and further verified by PCR using flanking primers. To generate a yeast expression construct for *SDO1*, the genomic sequence of *SDO1* (-200 to +200bp) was amplified by PCR from yeast genomic DNA using primers containing SpeI and EcoRI restriction sites. PCR amplification was performed using Phusion High-Fidelity PCR Master Mix (New England Biolabs). The amplified fragment was gel-purified and digested with SpeI and EcoRI, along with the yeast shuttle vector pRS-313. Following purification, the insert and vector were ligated using T4 DNA ligase. The ligation products were transformed into *Escherichia coli*, and positive clones were identified by PCR. Plasmid DNA was isolated from confirmed clones and used for yeast transformation. All constructs were verified by Sanger sequencing. Primers used were listed in the **Supplemental Table 4**. The pRS416 PTC1 plasmid was kindly provided by Dr. Lois S. Weisman (University of Michigan). Yeast transformations were performed using the lithium acetate method(*1*). Transformants were selected on synthetic complete (SC) medium lacking the appropriate nutrients. Human samples were obtained and analyzed in compliance with the Declaration of Helsinki, after written consent. All protocols were approved by the Ethics Committee of Azienda Ospedaliera Universitaria Integrata (Verona, Italy: approval No. 4182 CESC).

*Cell survival and cell cycle assays for HeLa cells*. HeLa cells ($1\times10^3$ cells per well in the 96-well plate) were exposed to various small molecules for 24, 48, and 72 h, respectively, until parental cells reached total cell death at higher dosage. We used the alamarBlue Cell Viability Reagent (ThermoFisher Scientific Inc., MA) to measure cell viability and proliferation. Cells were detached using TrypLE, fixed and permeabilized with -20°C chilled 70% ethanol on ice for 30 min. Fixed cells were washed twice with 1x PBS, incubated with 500 µL PBS containing 100 µg/mL RNase A (ThermoFisher Scientific Inc., MA) at room temperature for 30 min. Add 5 µL propidium iodide solution (PI 1 mg/mL, Sigma-Aldrich, MO). DNA content was assessed by BD FACSymphony A5 SE (Becton, Dickinson and Company, NJ) and the cell cycle analysis was performed using the Dean-Jett Fox model of FlowJo software (Becton, Dickinson and Company, NJ). For yeast cells, cells growing in mid-logarithmic phase were harvested by a brief centrifugation at 13000rpm for 1 min, washed with 1x PBS, and fixed in 70% ethanol overnight at 4ºC. Cells were centrifuged at the same speed to remove ethanol, and sterile distilled water was added to rehydrate the cells. After a brief centrifugation, RNase (0.25 mg/mL) was added to the samples and cells were incubated at 50°C for 2 h followed by incubation in 0.2 mg/mL of proteinase K for 2 h at 50°C in PBS. Samples were centrifuged and PI was added to the final concentration of 10 µg/mL. Sonicate the samples for 10 seconds and 0.25 OD600 equivalent cells were used to analysis the cellular DNA content using BD LSR Fortessa (Becton, Dickinson and Company, NJ).

*Growth curve assay for yeast cells.* Exponentially growing yeast cells were harvested and washed with sterile water. A 100 mL of 0.2 OD600 equivalent cells in YPD media was added in triplicate to wells in 96 well round bottom plates, and each well was overlaid with optical adhesive film to prevent evaporation. The plate was monitored using BioTek Synergy H1 Hybrid Multi-Mode Microplate Reader (Agilent Technologies, Inc., CA), taking A600 measurements at 1.5 h intervals for 7.5 h, with linear shaking occurring between readings at 30°C. The mean of the A600 reading for the triplicate strain was normalized to that of blank having YPD medium and plot the graph in the excel sheet.

*Drop-dilution assay.* Exponentially growing yeast cells were diluted to $OD_{600}$ 1. Three μL of cell culture starting at $OD_{600}$ 1 and serially diluted 5-fold were spotted onto YPD plates using a multichannel pipette. Spotted cells were allowed to grow at 30°C until good-sized colonies could be seen. Plates were imaged using a scanner.

*CRISPR editing and siRNA silencing.* The Alt-R CRISPR-Cas9 System was used to introduce indels in the HeLa to target *SBDS* exon 2, where hot spots exist in SDS patients. HeLa cells are known to have no mutations in *SBDS*. To form gRNA, crRNA targeting *SBDS* and tracrRNA were annealed, then S.p. Cas9 nuclease V3 protein was added to form ribonucleoprotein (Integrated DNA Technologies, Inc., IA). Cas9 ribonucleoprotein was introduced into parental HeLa cells using Lipofectamine CRISPRMAX (ThermoFisher Scientific Inc., MA) according to the manufacturer's protocol. Single cell sorting into 96-well plate was performed using FACSMelody (Becton, Dickinson and Company, NJ). Cells were screened by Western blotting for SBDS. Small interfering RNA duplexes (siRNA) against human *MAPK14* (encoding p38α), human *MAP3K20* (encoding ZAK) and the negative control pool were obtained from Dharmacon (Lafayette, CO). siRNA consisting of a mixture of 4 different RNA duplexes in equimolar concentrations was used for knockdown of *MAPK14* (ON-TARGETplus Human MAPK14 (1432) siRNA SMARTpool). For knockdown of ZAK, siRNA J-005068-13 was used which was selected by testing in HeLa cells to suppress most from set of 4 siRNAs (ON-TARGETplus Human MAP3K20 (51776) siRNA). HeLa cells were transfected with either specific siRNA or the control scrambled siRNA (ON-TARGETplus Non-targeting Control Pool) using the Lipofectamine RNAiMAX (ThermoFisher Scientific Inc., MA).

*Immunoprecipitation and immunoblotting*. For Western blotting of mammalian cells, cells were lysed in Pierce RIPA Buffer (ThermoFisher Scientific Inc., MA) supplemented with the protein and phosphatase inhibitor cocktail. The lysate was mixed with 4x Laemmli Sample Buffer (Bio-Rad Laboratories, CA) and denatured at 95°C for 5 min. For yeast cells, exponentially growing equivalent number of cells were harvested by brief centrifugation at 13000 rpm for 1 min. Total cellular protein was prepared by incubating cells with 0.2N NaOH for 10 min at room temperature. Following a brief centrifugation, the cell pellet was resuspended in 1x Laemmli sample buffer (125 mM Tris-HCl pH 6.8, 5% SDS, 0.004% Bromophenol Blue, 1.427M (5%) β-mercaptoethanol, 20% glycerol). Vortex the samples vigorously for 8–10 min and boiled at 99°C for 5 min. For immunoprecipitation, cell lysates were prepared using Pierce IP Lysis Buffer (ThermoFisher Scientific Inc., MA) supplemented with the protein and phosphatase inhibitor cocktail. Immunoprecipitation was performed using Dynabeads Protein A and G Immunoprecipitation Kit for rabbit and mouse antibodies (ThermoFisher Scientific Inc., MA), respectively, according to the manufacturer's protocol. The antibodies used for immunoprecipitation and immunoblotting are listed in the **Supplemental Table 1**. Signals were quantified using the ImageJ software (NIH, MD) and the ImageQuant TL analysis software (Cytiva, MA).

*RNA extraction and quantitative real-time PCR.* For mammalian cells, total RNA was isolated from the harvested cells using TRIzol reagent (ThermoFisher Scientific Inc., MA). Complementary DNA was synthesized using the iScript cDNA Synthesis Kit (Bio-Rad Laboratories, CA). For yeast cells, exponentially growing cells were treated with 0.8M NaCl. Total RNA was isolated at the indicated time points (0, 30 min) using the Qiagen RNeasy Mini Kit (QIAGEN Sciences, MD). 2μg of total RNA was reverse transcribed to synthesize complementary DNA using the High-Capacity RNA-to-cDNA Kit (ThermoFisher Scientific Inc., MA) according to the manufacturer's instructions. Quantitative real-time quantitative PCR was performed using PowerUp SYBR Green Master Mix (ThermoFisher Scientific Inc., MA). Relative gene expression was evaluated by the $2^{-\Delta\Delta CT}$ method. Primers used were listed in the **Supplemental Table 4**.

*Polysome Profiling and Ribo-Seq.* The appropriate volume of yeast lysate containing equal OD was treated with Turbo DNAse I and incubated at 25°C for 30 min. The cell extract is then loaded onto a 15%-45% sucrose gradient to separate the polysomes by ultra-centrifugation at 38k RPM for 2 hr at 4°C. The appropriate volume of yeast lysate containing equal OD was treated with Rnase I, and Turbo DNAse I and incubated at 25°C for 30 min; the digestion was stopped with SUPERase-inhibitor and placed on ice. The cell extract was then loaded onto a 15%-45% sucrose gradient to separate the polysomes by ultra-centrifugation at 41k RPM for 2 hours and 26 minutes at 4°C. The relevant fractions corresponding to the 80S monosome were collected and precipitated using ethanol, and the RNA was isolated by phenol-chloroform extraction. In parallel, total RNA was isolated from each sample and subjected to the Turbo DNase I digestion and then RNA-seq libraries were constructed using Next Ultra II directional RNA-seq kit (New England Biolabs). The ribosome-protected fragments were subjected to denaturing polyacrylamide-mediated gel electrophoresis and were gel-extracted based on size 25 nt to 35 nt. The RNA was precipitated and subjected to the RiboMinus Ribosomal Eukaryotic RNA depletion kit (Thermo Fisher). The rRNA-depleted RNA footprints were then subjected to de-phosphorylation. Adapter ligation, reverse transcription and subsequent cDNA amplification was performed using the NEXT-Flex smRNA-Seq Kit v4 (PerkinElmer); cDNA library QC was performed by Tapestation before sequencing on an Illumina NovaSeq X single-end for 50 cycles.

*Ribosome Profiling Data Processing and Stalling Analysis.* Ribosome profiling reads in fastq format (Single-end sequencing) were first processed using FastQC (v0.12.1) for QC analysis. The reads were then aligned to saccharomyces cerevisiae S288C (sacCer3; saccharomyces_cerevisiae_R64-1-1 sourced on Saccharomyces Genome Database (SGD)) using STAR (v2.5.2b), and the unique mapping rates for bulk RNA were between 51% to 66%. For RPF data, the pre-process steps were applied. The fastq_quality_filter module was used to extract the bases with 99% accuracy based on Q Score in fastq and the reads where less than 70% of bases with an accuracy for at least 99% were removed. Module fastx_collapser (FASTX Toolkit 0.0.14) was used to perform sequence collapse and Cutadapt (v4.6) was applied to remove universal Illumina adaptor. The mapping was then performed on the re-preprocessed RPF data and the unique mapped rates were between 13% to 33%. Since the reference is built on open reading frame (ORF) regions, the ORF based gene count table for RNA and RPF were the input for the downstream DE analysis (DESeq2 (v1.42.0)) and TE analysis. For ribosome stalling analysis, mapped results were first filtered to retain read length between 26 and 34 nucleotides, with expected peaks between 28 and 30 nt. Then, processed mapped results were analyzed using the ORFik (v4.1) R package following the yeast sequencing data analysis process provided in the R package. Transcript and gene annotations were derived from Ensembl R64 GTF files and supplemented with a custom annotation table for 5' and 3' UTR coordinates to account for the regions that are not annotated in standard genomic files.

*Stalling Metrics and Differential Analysis.* Ribosome stalling was quantified using a stalling score (***S***) and the coefficient of variation (***CV***) to measure coverage uniformity across transcripts. The stalling score was calculated as:

$$\boldsymbol{S} = max(C_i)/(C_{mean}+1)$$

where $C_i$ represents the coverage at position $i$ and $C_{mean}$ represents the mean coverage. Peak coverage is calculated by finding the maximum coverage $max(C_i)$ within a given transcript. A pseudo-count of 1 was added to $C_{mean}$ to prevent division by zero. Coverage variation was measured as:

$$\boldsymbol{CV} = \sigma / (C_{mean} + 0.1)$$

where $\sigma$ represents the standard deviation of coverage across the region. A pseudo-count of 0.1 was added to $C_{mean}$ to prevent division by zero. All analysis was restricted to high-confidence genes containing data in ≥ 2 replicates per condition (WT and *sdo1Δ*; total 4663 genes). Differential occupancy was determined by calculating the $\log_2$ fold change of the mean scores:

$$\boldsymbol{log_2FC} = log_2((C_{sdo1\Delta} + 1) / (C_{WT}+1))$$

Statistical significance between WT and *sdo1D* stalling score distributions was assessed using Welch's t-test, while the Kolmogorov-Smirnov (KS) test was employed to identify significant shifts in cumulative distribution functions (ECDF).

*Stress Granule Immunofluorescence and Quantification.* Wild-type (WT) and *SBDS−/−* (KO) HeLa cells were seeded in black-walled, clear-bottomed 96-well imaging plates (Costar, #3904) at a density of 8×10^3 cells per well. Wells were pre-coated with collagen (Sigma, #C8919), diluted 1:50 in water, incubated at 37C for 30min, then aspirated prior to cell seeding. Following cell attachment, stress granule formation was induced by treating cells with varying concentrations of sodium arsenite (From 1μM to 1mM) for 45 minutes at 37°C. Cells were fixed in 4% paraformaldehyde (PFA) for 10 minutes at room temperature, followed by a wash with phosphate-buffered saline (PBS). For immunostaining, cells were permeabilized with 0.1% Triton X-100 in PBS for 10 minutes at RT with gentle agitation. After washing twice with PBS containing 0.1% Tween-20 (PBST), non-specific binding was blocked using a blocking buffer (2% BSA in PBST) for 1 hour at RT. Cells were then incubated with a primary antibody against the stress granule marker G3BP1 (Sigma, Cat# G6046; 1:1000 dilution) in PBST overnight at 4°C. Following two washes with PBST, cells were incubated with an anti-mouse secondary antibody conjugated to Alexa Fluor 594 (ThermoFisher, #A-11032; 1:1000) for 30 minutes at RT, protected from light. Nuclei were counterstained with DAPI (1:1000 in PBST) for 5 minutes. After two final washes with PBST, plates were stored in PBST at 4°C until imaging.

*Imaging and Data Analysis.* Plates were imaged using a Cytation 7 Cell Imaging Multimode Reader (BioTek) at 20x magnification. Quantitative analysis of stress granules was performed using CellProfiler (v.4.2.4) according to a custom pipeline, Briefly, nuclei were segmented using the DAPI channel, and cell boundaries were defined using the IdentifySecondaryObjects module. G3BP1-positive stress granules were identified within cells using the IdentifySecondaryObjects module set to 10-30 pixel object widths. Further thresholding and filtering were applied to remove membrane artifacts. In all cases, positive controls (arsenite-treated WT cells) or negative controls (WT cells alone) were used to benchmark SG-positive cell calling. Data visualization was conducted using GraphPad Prism.

*Protein Synthesis Assay*. The assay was performed using Click-iT Plus OPP Alexa Fluor 647 Protein Synthesis Assay Kit (Thermo Fisher Scientific Inc., MA) with minor modifications described elsewhere.(*2, 3*) Briefly, cells were treated with O-propargyl-puromycin (OPP), a puromycin analogue that is incorporated in nascent proteins, for 30 min in full media. For negative controls, cells were treated with cycloheximide (CHX) from 15 min prior to and during OPP treatment. Cells were detached using TrypLE, fixed and permeabilized with -20°C chilled 70% ethanol on ice for 30 min. Incorporated OPP was labeled with Alexa Fluor 647 by click reaction, followed by stain with DAPI. Fluorescent intensity was assessed by FACSymphony A5 SE (Becton, Dickinson and Company, NJ).

*Statistical analysis.* Statistical analysis was conducted using GraphPad Prism 10.1.2 (GraphPad Software Inc., MA). A two-way ANOVA with Tukey's multiple comparisons was used to estimate the mean differences between groups. A $P$ value of $<.05$ was considered statistically significant.

**References**

1. R. D. Gietz, R. H. Schiestl, Large-scale high-efficiency yeast transformation using the LiAc/SS carrier DNA/PEG method. *Nat Protoc* **2**, 38-41 (2007).
2. J. C. Hsu, J. B. Pawlak, M. Laurent-Rolle, P. Cresswell, Protocol for assessing translational regulation in mammalian cell lines by OP-Puro labeling. *STAR Protoc* **3**, 101654 (2022).
3. L. Hidalgo San Jose, R. A. J. Signer, Cell-type-specific quantification of protein synthesis in vivo. *Nat Protoc* **14**, 441-460 (2019).

**Fig. S1. Differential growth in WT versus s*do1Δ* yeast.**

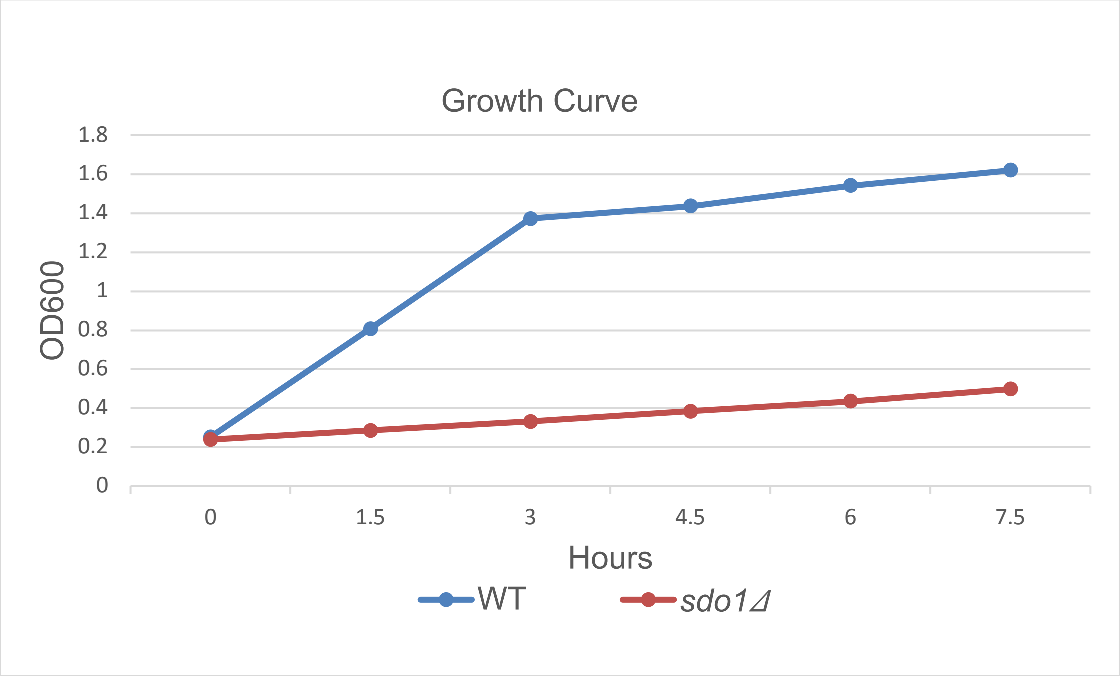

**Fig. S2. CRISPR-Cas9-mediated gene editing in HeLa cells.**

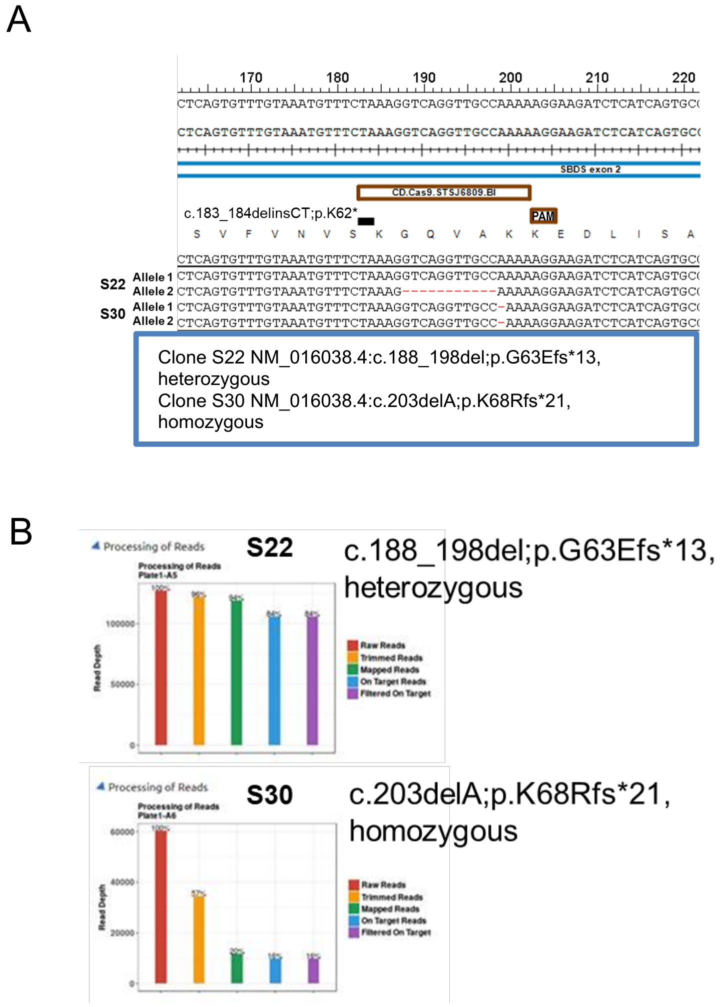


(**A**) Schematic representation of CRISPR-Cas9 targeting exon 2 of *SBDS*. (**B**) Allele frequencies of individual clones determined by amplicon deep sequencing of *SBDS* exon 2.

**Fig. S3. Protein and phosphorylation levels in SBDS-WT and SBDS-KO HeLa cells following hydrogen peroxide exposure.**

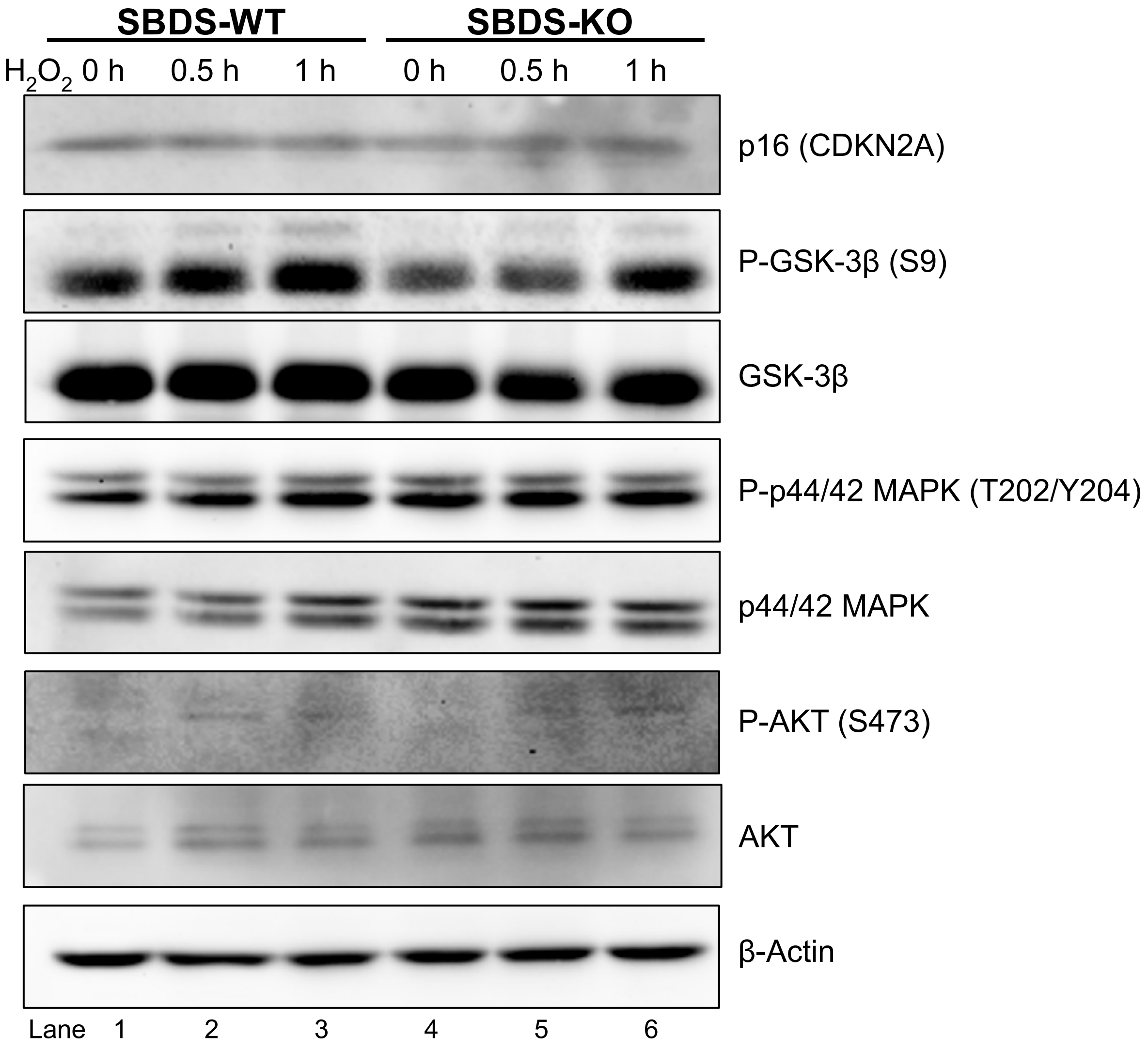

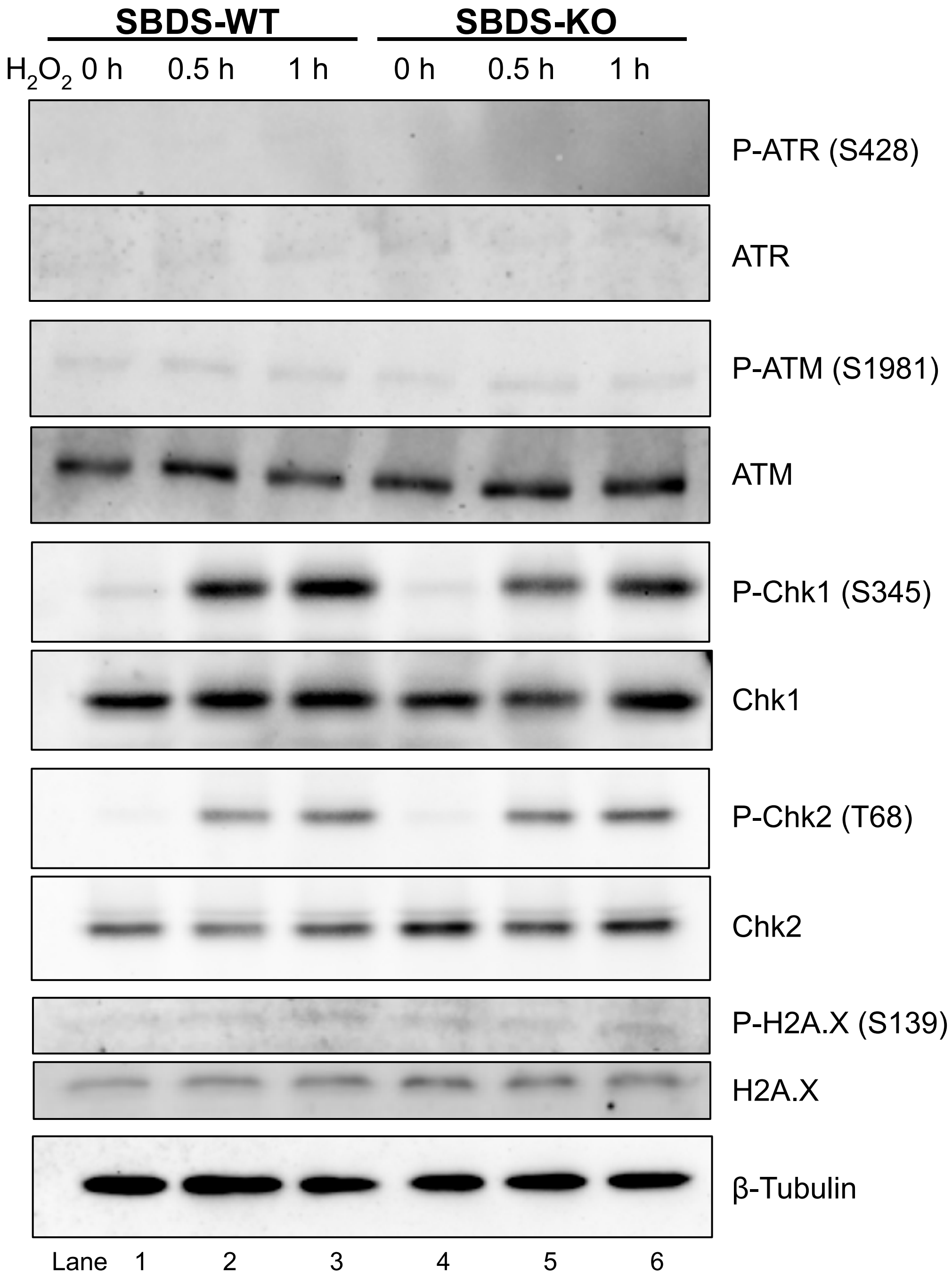


Levels of p16, phospho-GSK-3β, phospho-p44/42 MAPK, and γH2A.X did not differ between *SBDS* genotypes. AKT and ATM exhibited slight phosphorylation upon hydrogen peroxide ($H_2O_2$) stimulation, whereas CHK1 and CHK2 were significantly phosphorylated in response to $H_2O_2$. These phosphorylation levels were not significantly different between *SBDS* genotypes.

**Fig. S4. Phosphorylation of p38 SAPK following hydrogen peroxide exposure in HeLa cells.**

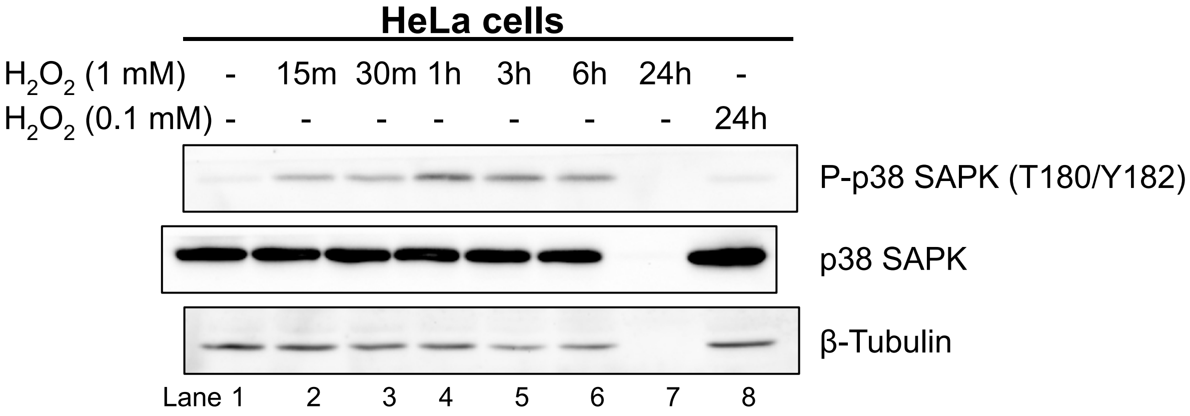


HeLa parental cells were treated with 1 mM or 0.1 mM hydrogen peroxide ($H_2O_2$) for the indicated times. Phosphorylation of p38 SAPK peaked after 1 h of incubation. Treatment with 1 mM $H_2O_2$ for 24 h induced complete cell death, resulting in protein depletion (lane 7), whereas 0.1 mM $H_2O_2$ did not increase p38 phosphorylation (lane 8).

**Fig. S5. Overexpression of Ptc1 yeast decreased osmotic stress-induction of phosphorylation of Hog1.** Yeast were treated with 0.8 M NaCl for 30 min at RT. Lysates were then separated b SDS-PAGE and transferred onto nitrocellulose and probed with anti-phospho-HOG1 antibody. The blot was afterward stripped and reprobed anti-HOG1 antibody.

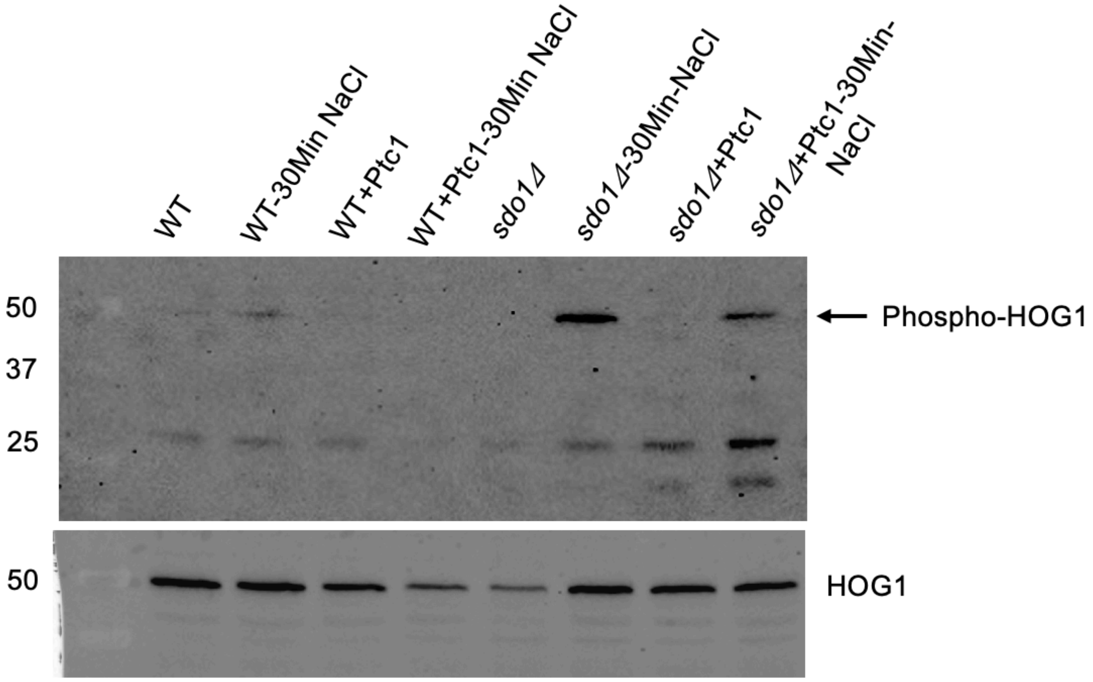

**Fig. S6. Dose–response curve of SBDS-WT versus SBDS-KO HeLa cells after VX-745 treatment.**

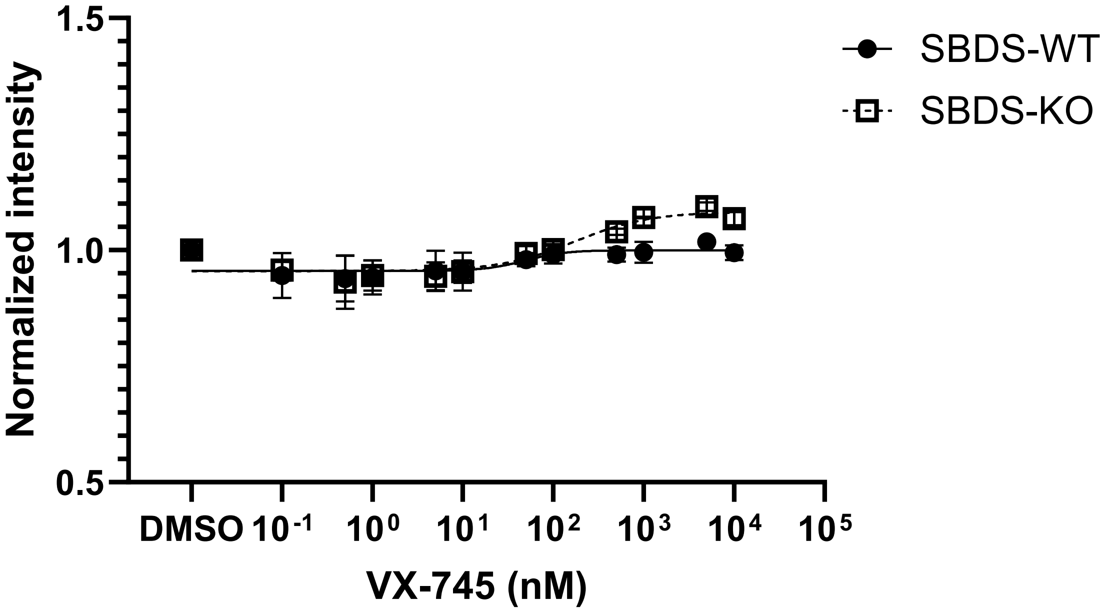


HeLa cells were treated with various concentrations of the potent p38α inhibitor VX-745 for 7 days. Cell survival was assessed using the alamarBlue assay.

**Fig. S7. Efficiency of gene silencing by siRNA targeting *MAPK14* and *MAP3K20* in HeLa cells.**

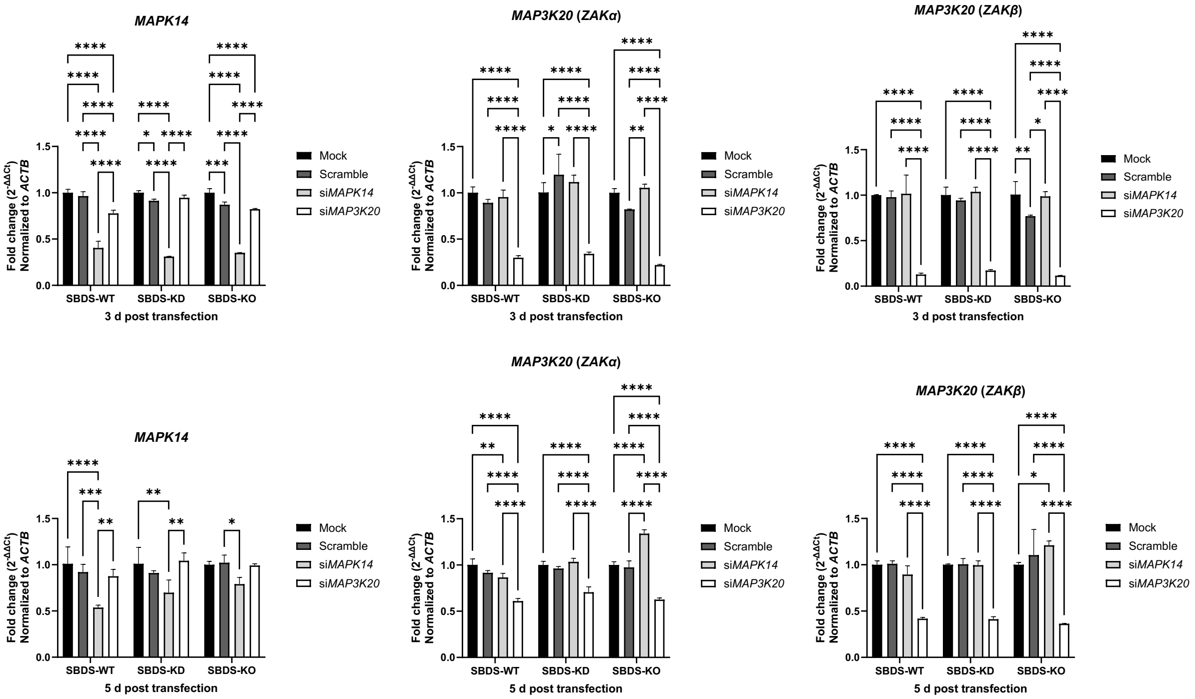


Gene silencing profiles were assessed at 3 and 5 days following transfection with siRNA targeting *MAPK14* and *MAP3K20*, respectively. Gene expression was normalized to *ACTB* and expressed as the relative fold change to the mock-treated control group. To distinguish alternative splice variants encoding different isoforms (ZAKα and ZAKβ), variant-specific primers were used to detect *MAP3K20*. Each value represents the mean ± SD (n = 3). * $P<0.05$, ** $P<0.01$, *** $P<0.001$, **** $P<0.001$; two-way ANOVA with Tukey's correction.

**Fig. S8. Reactive oxygen species scavengers did not improve cell survival in SBDS-deficient HeLa cells.**

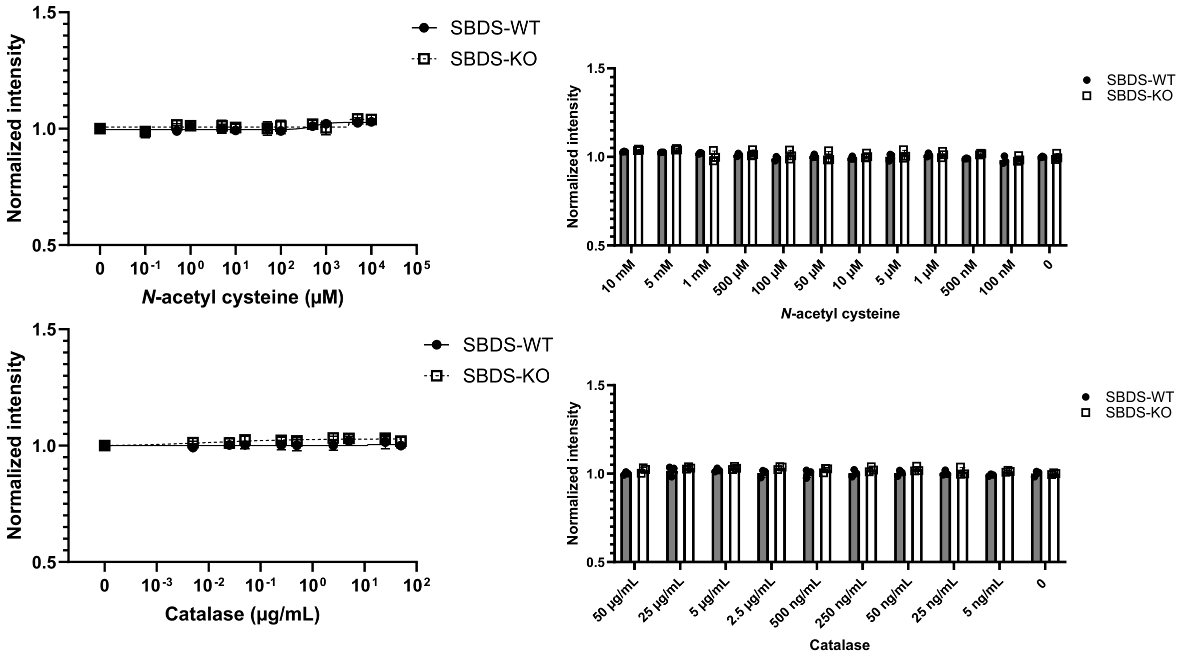


Cell survival, assessed using the alamarBlue assay, showed no significant change after treatment with catalase (from bovine liver) or *N*-acetylcysteine in either SBDS-WT or SBDS-KO cells.

**Fig. S9. Reactive oxygen species scavengers did not decrease spontaneous p38 phosphorylation.**

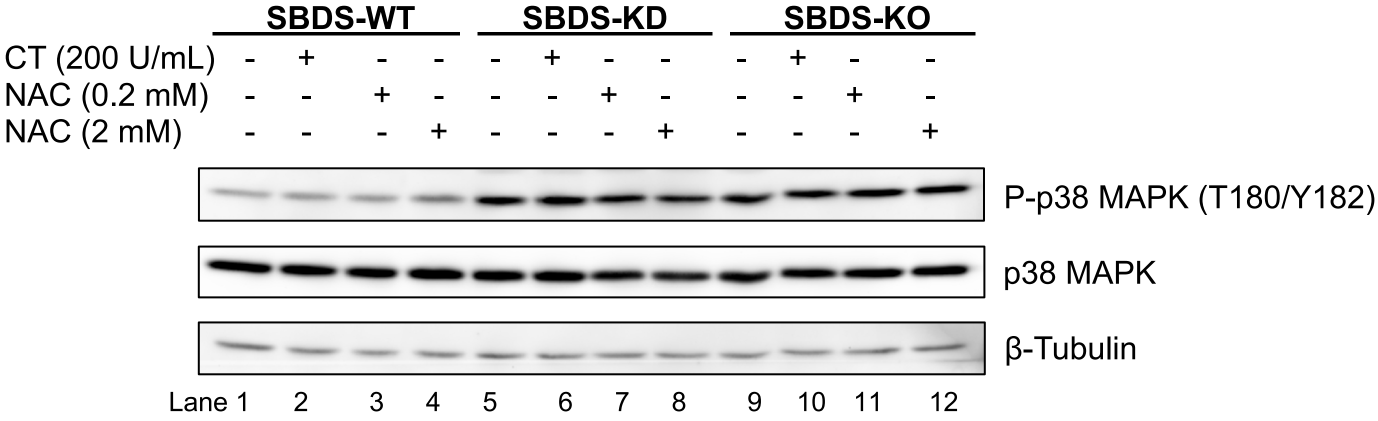


SBDS-WT, SBDS-KD, and SBDS-KO cells were treated with either catalase (CT, from bovine liver) or *N*-acetylcysteine (NAC) for 24 h. These treatments did not alter spontaneous phosphorylation levels of p38 in SBDS-KD or SBDS-KO cells.

**Fig. S10. Hydrogen peroxide did not induce significant threonine phosphorylation of ZAKα in HeLa cells.**

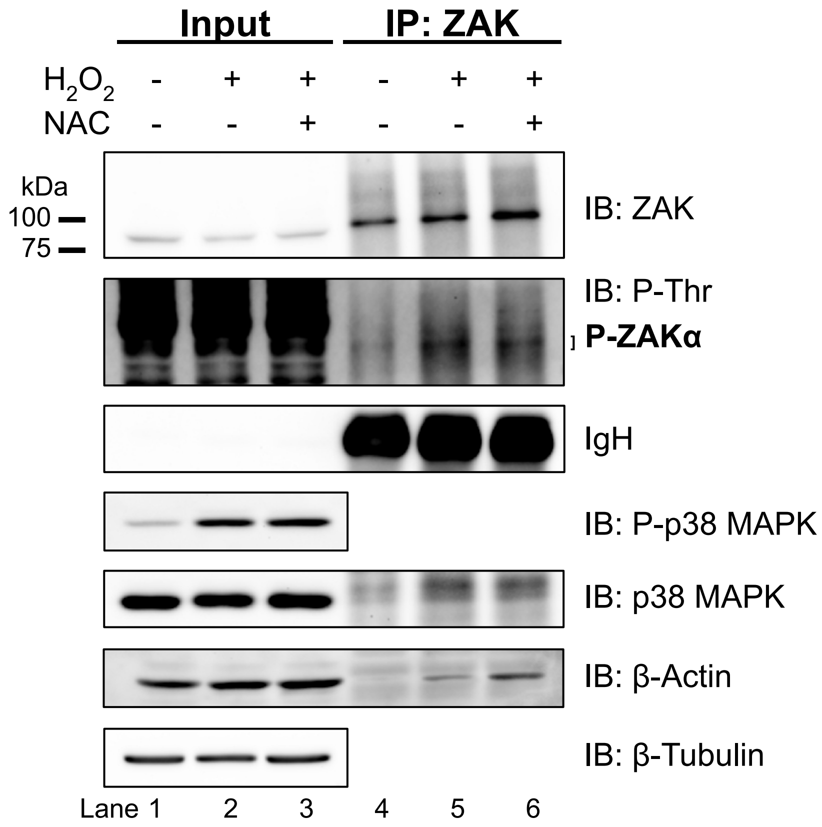


HeLa parental cells were treated with hydrogen peroxide (1 mM $H_2O_2$) for 0.5 h, which induced phosphorylation of p38 SAPK. Pretreatment with N-acetyl-L-cysteine (2 mM NAC) for 24 h prior to $H_2O_2$ exposure did not significantly mitigate p38 phosphorylation. Immunoprecipitation (IP) using anti-ZAK antibodies followed by immunoblotting (IB) revealed no significant difference in ZAKα phosphorylation regardless of $H_2O_2$ or NAC treatment.

**Fig. S11. Antioxidative agents did not alleviate spontaneous phosphorylation of ZAKα in SBDS-deficient HeLa cells.**

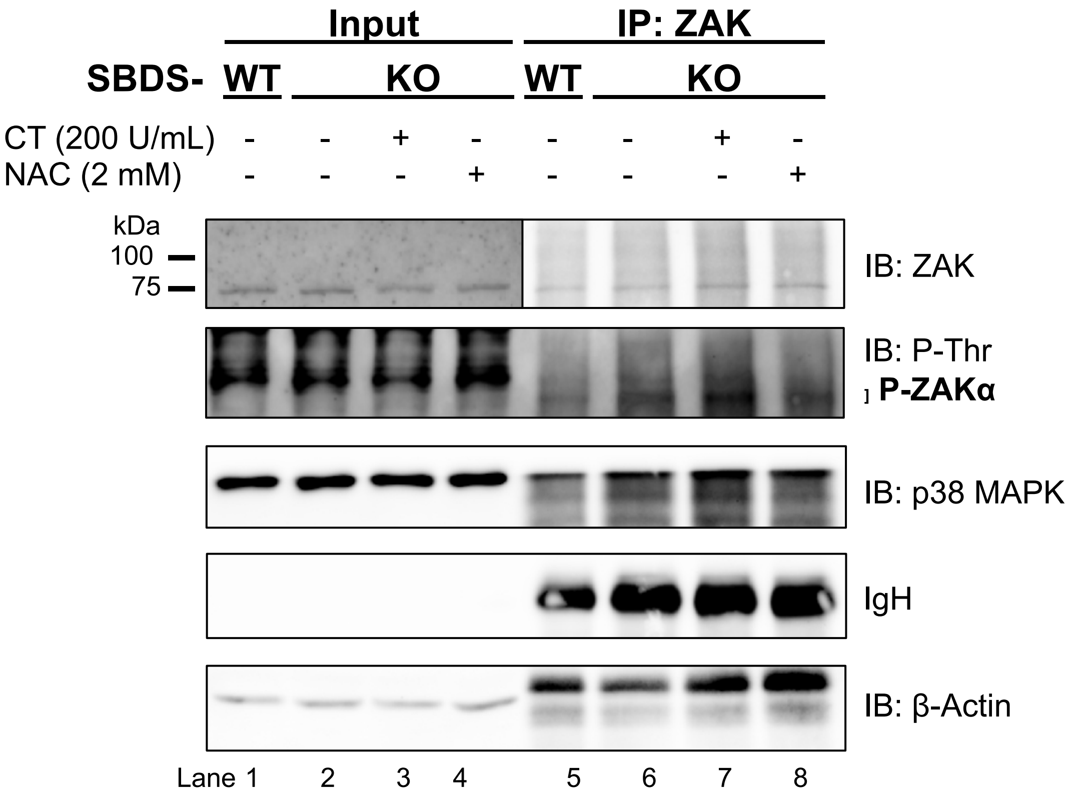


Immunoprecipitation (IP) using anti-ZAK antibodies followed by immunoblotting (IB) was performed in SBDS-KO cells treated with either catalase (CT, from bovine liver) or *N*-acetyl-L-cysteine (NAC) for 24 h. ZAKα was more phosphorylated in untreated SBDS-KO cells compared with SBDS-WT cells. Increased ZAKα phosphorylation in untreated SBDS-KO cells promoted greater binding of downstream p38 compared with SBDS-WT cells. Phosphorylation levels of ZAKα and precipitated p38 protein levels were comparable among SBDS-KO cells regardless of antioxidant treatment.

**Table S1. Patient characteristics.**

| UPN | Sex | Age | *SBDS* mutation | ANC (/µL) | Phenotype | MDS/ AML |
|---|---|---|---|---|---|---|
| **63** | M | 15 | c.258+2T>C<br>c.183_184TA>CT | 1290 | RPI, bone malformations | No |
| **75** | F | 10 | c.258+2T>C<br>c.183_184TA>CT | 920 | EPI, thrombocytopenia, bone malformations | No |
| **94** | F | 19 | c.258+2T>C<br>c.183-184TA>CT | 1750 | EPI, bone malformations | No |

Abbreviations: UPN, unique patient number; M, male; F, female; ANC, absolute neutrophil count; EPI, exocrine pancreatic insufficiency; MDS, myelodysplastic syndrome; AML, acute myelogenous leukemia.

**Table S2. Antibodies used in this study.**

| REAGENT or RESOURCES | SOURCE | IDENTIFIER |
|---|---|---|
| SBDS (D-6) Mouse mAb | Santa Cruz Biotechnology | sc-271600; RRID:AB_10650682 |
| EFL1 (RIA1) (A-7) Mouse mAb | Santa Cruz Biotechnology | sc-514617 |
| eIF6 Rabbit pAb | Cell signaling technology | CST#3263; RRID:AB_2293295 |
| p53 (DO-1) Mouse mAb | Santa Cruz Biotechnology | sc-126; RRID:AB_628082 |
| p21 (Waf1/Cip1, CDKN1A) (12D1) Rabbit mAb | Cell signaling technology | CST#2947; RRID:AB_823586 |
| Phospho-p38 MAPK (Thr180/Tyr182) Rabbit pAb | Cell signaling technology | CST#9211; RRID:AB_331641 |
| p38 MAPK (D13E1) XP Rabbit mAb | Cell signaling technology | CST#8690; RRID:AB_10999090 |
| Phospho-SAPK/JNK (Thr183/Tyr185) (81E11) Rabbit mAb | Cell signaling technology | CST#4668; RRID:AB_823588 |
| SAPK/JNK (D-2) Mouse mAb | Santa Cruz Biotechnology | sc-7345; RRID:AB_675864 |
| Phospho-MAPKAPK-2 (Thr334) (27B7) Rabbit mAb | Cell signaling technology | CST#3007; RRID:AB_490936 |
| Phospho-MAPKAPK-2 (Thr222) (9A7) Rabbit mAb | Cell signaling technology | CST#3316; RRID:AB_2141311 |
| MAPKAPK-2 (D1E11) Rabbit mAb | Cell signaling technology | CST#12155; RRID:AB_2797831 |
| ZAK Rabbit pAb | Fortis Life Sciences (Bethyl Laboratories) | A301-993A; RRID:AB_1576612 |
| Phospho-Threonine (42H4) Mouse mAb | Cell signaling technology | CST#9386; RRID:AB_331239 |
| p16 INK4A (D3W8G) Rabbit mAb | Cell signaling technology | CST#92803; RRID:AB_2750891 |
| Phospho-GSK-3β (Ser9) (5B3) Rabbit mAb | Cell signaling technology | CST#9323; RRID:AB_2115201 |
| GSK-3β (27C10) Rabbit mAb | Cell signaling technology | CST#9315; RRID:AB_490890 |
| Phospho-p44/42 MAPK (ERK1/2) (Thr202/Tyr204) Rabbit pAb | Cell signaling technology | CST#9101; RRID:AB_331646 |
| p44/42 MAPK (ERK1/2) Rabbit pAb | Cell signaling technology | CST#9102; RRID:AB_330744 |
| Phospho-Akt (Ser473) (D9E) XP Rabbit mAb | Cell signaling technology | CST#4060; RRID:AB_2315049 |

| | | |
|---|---|---|
| Akt (pan) (40D4) Mouse mAb | Cell signaling technology | CST#2920; RRID:AB_1147620 |
| Phospho-ATM (Ser1981) (D6H9) Rabbit mAb | Cell signaling technology | CST#5883; RRID:AB_10835213 |
| ATM (D2E2) Rabbit mAb | Cell signaling technology | CST#2873; RRID:AB_2062659 |
| Phospho-ATR (Ser428) Rabbit pAb | Cell signaling technology | CST#2853; RRID:AB_2290281 |
| ATR Rabbit pAb | Cell signaling technology | CST#2790; RRID:AB_2227860 |
| Phospho-Chk1 (Ser345) (133D3) Rabbit mAb | Cell signaling technology | CST#2348; RRID:AB_331212 |
| Chk1 (2G1D5) Mouse mAb | Cell signaling technology | CST#2360; RRID:AB_2080320 |
| Phospho-Chk2 (Thr68) (C13C1) Rabbit mAb | Cell signaling technology | CST#2197; RRID:AB_2080501 |
| Chk2 (D9C6) Rabbit mAb | Cell signaling technology | CST#6334; RRID:AB_11178526 |
| Phospho-Histone H2A.X (Ser139) (20E3) Rabbit mAb | Cell signaling technology | CST#9718; RRID:AB_2118009 |
| Histone H2A.X (D17A3) XP Rabbit mAb | Cell signaling technology | CST#7631; RRID:AB_10860771 |
| Phospho-eIF2α (Ser51) (D9G8) XP Rabbit mAb | Cell signaling technology | CST#3398; RRID:AB_2096481 |
| eIF2α Rabbit pAb | Cell signaling technology | CST#9722; RRID:AB_2230924 |
| β-Actin (C4) Mouse mAb | Santa Cruz Biotechnology | sc-47778; RRID:AB_626632 |
| β-Tubulin (9F3) Rabbit mAb | Cell signaling technology | CST#2128; RRID:AB_823664 |
| Rabbit IgG (H&L) Peroxidase Conjugated Donkey pAb | Rockland Immunochemicals | 611-7302; RRID:AB_219747 |
| Mouse IgG (H&L) Peroxidase Conjugated Sheep pAb | Rockland Immunochemicals | 610-603-002; RRID:AB_219694 |
| Hog1 Mouse mAb | Santa Cruz Biotechnology | Sc-165978 |
| Phospho-pHog1/p38 MAPK Rabbit mAb | Cell signaling technology | CST#9215S |

**Table S3. Chemicals, peptides, and recombinant proteins used in this study.**

| REAGENT or RESOURCES | SOURCE | IDENTIFIER |
|---|---|---|
| TrypLE Express Enzyme (1X), no phenol red | Thermo Fisher Scientific | 12604021 |
| Trypan Blue 0.4% Solution | Lonza Bioscience | 17942E |
| alamarBlue Cell Viability Reagent | Thermo Fisher Scientific | DAL1025 |
| Dynabeads Protein A Immunoprecipitation Kit | Thermo Fisher Scientific | 10006D |
| Dynabeads Protein G Immunoprecipitation Kit | Thermo Fisher Scientific | 10007D |
| Pierce RIPA Buffer | Thermo Fisher Scientific | 89900 |
| Pierce IP Lysis Buffer | Thermo Fisher Scientific | 87787 |
| Halt Protease Inhibitor Cocktail | Thermo Fisher Scientific | 78430 |
| Halt Phosphatase Inhibitor Cocktail | Thermo Fisher Scientific | 78420 |
| Sodium orthovanadate | Sigma-Aldrich | 450243-10G |
| 2-Mercaptoethanol | Sigma-Aldrich | M6250-100ML |
| 4x Laemmli Sample Buffer | Bio-Rad Laboratories | 1610747 |
| Tris Buffered Saline with 1% Casein | Bio-Rad Laboratories | 1610782 |
| Blotto Immunoanalytical Grade (Non-Fat Dry Milk) | Rockland Immunochemicals | B501-0500 |
| Amersham ECL start Western Blotting Detection Reagent | Cytiva | RPN3243 |
| Amersham ECL Prime Western Blotting Detection Reagent | Cytiva | RPN2232 |
| DNeasy Blood & Tissue Kit | Qiagen | 69504 |
| QIAquick PCR Purification Kit | Qiagen | 28104 |
| TRIzol Reagent | Thermo Fisher Scientific | 15596018 |
| Chloroform ≥99.8% stabilized EtOH | Acros Organics | 423555000 |
| iScript cDNA Synthesis Kit | Bio-Rad Laboratories | 1708890 |
| PowerUp SYBR Green Master Mix for qPCR | Thermo Fisher Scientific | A25777 |
| Mitomycin C | Sigma Aldrich | M5353-0.2ML |
| Anisomycin | Sigma-Aldrich | A5862-0.5ML |
| MG-132 | Sigma-Aldrich | M7449-200UL |
| CX-5461 (pidnarulex) | Selleck Chemicals | S2684 |
| Actinomycin D | Sigma-Aldrich | A9415-2MG |
| Menadione sodium bisulfite | Sigma-Aldrich | M5750-25G |
| VX-745 (neflamapimod) | Sigma-Aldrich | SML1638-5MG |
| Calyculin A | Sigma-Aldrich | 5.08226 |
| Olaparib | Sigma-Aldrich | SML3705 |

| | | |
|---|---|---|
| Daunorubicin hydrochloride | Sigma-Aldrich | 30450 |
| N-Acetyl-L-cysteine | Sigma-Aldrich | A7250-10G |
| Catalase from bovine liver | Sigma-Aldrich | C1345 |
| Alt-R CRISPR-Cas9 tracrRNA, ATTO 550 | Integrated DNA Technologies | 1075927 |
| Alt-R S.p. Cas9 Nuclease V3 | Integrated DNA Technologies | 1081058 |
| Lipofectamine CRISPRMAX Cas9 Transfection Reagent | Thermo Fisher Scientific | CMAX00001 |
| Lipofectamine RNAiMAX Transfection Reagent | Thermo Fisher Scientific | 13778075 |
| Click-iT Plus OPP Alexa Fluor 647 Protein Synthesis Assay Kit | Thermo Fisher Scientific | C10458 |

**Table S4. Oligonucleotides used in this study.**

| REAGENT or RESOURCES | SOURCE | IDENTIFIER |
|---|---|---|
| ON-TARGETplus Human MAPK14 (1432) siRNA SMARTpool | Dharmacon | L-003512-00-0005 |
| ON-TARGETplus siRNA Human MAPK14 #1 (GGAAUUCAAUGAUGUGUAU) | Dharmacon | J-003512-20 |
| ON-TARGETplus siRNA Human MAPK14 #2 (UCUCCGAGGUCUAAAGUAU) | Dharmacon | J-003512-21 |
| ON-TARGETplus siRNA Human MAPK14 #3 (GUAAUCUAGCUGUGAAUGA) | Dharmacon | J-003512-22 |
| ON-TARGETplus siRNA Human MAPK14 #4 (GUCCAUCAUUCAUGCGAAA) | Dharmacon | J-003512-23 |
| ON-TARGETplus Human MAP3K20 (51776) siRNA Set of 4 | Dharmacon | LQ-005068-00-0002 |
| ON-TARGETplus siRNA Human ZAK #1 (CGAGAAAGACGUUUAAAGA) | Dharmacon | J-005068-11 |
| ON-TARGETplus siRNA Human ZAK #2 (GGAUAUCACAGGACMGGA) | Dharmacon | J-005068-12 |
| ON-TARGETplus siRNA Human ZAK #3 (GUAAUCUAGCUGUGAAUGA) | Dharmacon | J-005068-13 |
| ON-TARGETplus siRNA Human ZAK #4 (GUCCAUCAUUCAUGCGAAA) | Dharmacon | J-005068-14 |
| ON-TARGETplus Non-targeting Control Pool | Dharmacon | D-001810-10-05 |
| CD.Cas9.STSJ6809.BI (Alt-R crRNA) (TAAAGGTCAGGTTGCCAAAA) | This paper | N/A |
| hSBDS Fwd2 Intron1 (CACATAGGACTTTCTCTCCTGCCC) | This paper | N/A |
| hSBDS Rev Intron2 (CTGAGTGCAGTGAAGTGCACTG) | This paper | N/A |
| hSBDS qPCR F (CAGATCCGCCTAACCAATGT) | This paper | N/A |
| hSBDS qPCR R (TGGGTCTGCAGAACTTCATC) | This paper | N/A |
| EFL1qPCRFwd (CTTCTGATGGGAAGGGAACTGG) | This paper | N/A |
| EFL1qPCRRev (ATGGCAGGCTACACAGTGTTGC) | This paper | N/A |
| Fwd hEIF6 qPCR 3156338 (CCGACCAGGTGCTAGTAGGAA) | This paper | N/A |
| Fwd hEIF6 qPCR 3156338 (CAGAAGGCACACCAGTCATTC) | This paper | N/A |
| HsPPP2CA_qPCR_Fwd (GGTGGTCTCTCGCCATCTATAG) | This paper | N/A |
| HsPPP2CA_qPCR_Rev (CTGGATCTGACCACAGCAAGTC) | This paper | N/A |
| HsMAPK14_qPCR_Fwd (GAGCGTTACCAGAACCTGTCTC) | This paper | N/A |

| | | |
|---|---|---|
| HsMAPK14_qPCR_Rev (AGTAACCGCAGTTCTCTGTAGGT) | This paper | N/A |
| HsMAP3K20_var1_F (GCAGTCCAACTTGCCATTCAGAC) | This paper | N/A |
| HsMAP3K20_var1_R (GCAGTCCAACTTGCCATTCAGAC) | This paper | N/A |
| HsMAP3K20_var2_F (CAGGAGCTGTGATGCATTCT) | This paper | N/A |
| HsMAP3K20_var2_R (CCAGAGCCATGTTGACTTTCT) | This paper | N/A |
| qPCR hActinb Fwd (CCTGTACGCCAACACAGTGC) | This paper | N/A |
| qPCR hActinb Rev (CTTCATTGTGCTGGGTGCCAG) | This paper | N/A |
| SDO1A Fwd (GAACAAACGAAATCTAAAAGCAAAA) | This paper | N/A |
| SDO1B Rev (CTTCTCAATAAGTTCGGGTTGTCTA) | This paper | N/A |
| SDO1C Fwd (CAAGACTATCGAAAAGGAATTGAAA) | This paper | N/A |
| SDO1D Rev (GACGATGAGGAAGAAGAATTTGTAA) | This paper | N/A |
| KanB Rev (CTGCAGCGAGGAGCCGTAAT) | This paper | N/A |
| KanC Fwd (TGATTTTGATGACGAGCGTAAT) | This paper | N/A |
| qPCR GPD1 Fwd (GACGCCTGGGAATGTGAAAA) | This paper | N/A |
| qPCR GPD1 Rev (TCTTCGACAGAGCCACATGT) | This paper | N/A |
| Sdo1 Fwd (AAGTGAATTCTATGTATTGACTGTTATTTA) | This paper | N/A |
| Sdo1 Rev (GCAGACTAGTGAACTATATAGGAGAGTTAT) | This paper | N/A |
